# A single-cell multi-omic and spatial atlas of B-cell lymphomas reveals differentiation drives intratumor heterogeneity

**DOI:** 10.1101/2023.11.06.565756

**Authors:** Donnacha Fitzgerald, Tobias Roider, Marc-Andrea Baertsch, Artur Kibler, Anastasiia Horlova, Erin Chung, Harald Vöhringer, Anna Mathioudaki, Bettina Budeus, Verena Passerini, Mareike Knoll, Johannes Mammen, Linsha Li, Léandra Caillé, Felix Czernilofsky, Peter-Martin Bruch, Nora Liebers, Matthias Meyer-Bender, Reiner Siebert, Oliver Weigert, Judith Zaugg, Garry Nolan, Marc Seifert, Sascha Dietrich, Wolfgang Huber

## Abstract

Intratumor heterogeneity underpins cancer pathogenesis and evolution, although it is typically considered independent from the differentiation processes that drive physiological cell-type diversity. As cancer types and subtypes arise from different cell types, we investigated whether cellular differentiation influences intratumor heterogeneity. Nodal B-cell non-Hodgkin lymphomas are a diverse set of cancers originating from different stages of B-cell maturation. Through single-cell transcriptome and surface epitope profiling (CITE-Seq) of diffuse large B-cell, mantle cell, follicular, and marginal zone lymphomas in addition to reactive lymph nodes from 51 patients, we found multiple B-cell maturation states within tumors. Intratumor maturation states emerged from the same clone, revealing divergent differentiation from a shared cell of origin. Maturation state composition varied across subtypes and tumors, which encompassed mixed cell-of-origin diagnostic subtypes. Through highly multiplexed immunohistochemistry (CODEX) of samples from 19 of these patients, we found that intratumor maturation states inhabited distinct spatial niches, displaying cellular interactions and regulatory networks typical of their maturation states while harboring different genetic variants. By deconvoluting intratumor maturation states from a microarray dataset of 507 patients, we identified risk groups within diagnoses with striking differences in survival, including IgM memory-enriched germinal center B-cell (M = 1.9 vs >10 years, p = 0.00039) and activated B-cell (M = 2.4 vs 9.6 years, p = 0.016) diffuse large B-cell lymphoma, and dark zone-enriched follicular lymphoma (M = 8.6 vs 13 years; p = 0.0019). Our findings reveal cellular differentiation remains plastic in B-cell lymphomas, driving tumor variation, evolution, and response.

**Key Points:**

- Cellular differentiation remains plastic in B-cell lymphomas, driving tumor variation, evolution, and response.
- Intratumor maturation states occupy unique immune niches, harbor distinct genetic variants, and are tied to different survival outcomes.

## Introduction

Cellular differentiation gives rise to the diversity of cell types and states observed across organs, tissues, and developmental lineages. Distinct cancer types and subtypes can emerge from these. The cell of origin, defined as the cell that acquired the first genetic aberrations that then lead to cancer, is widely thought to be a major determinant of tumor behavior and response to therapy^1^. The molecular and functional features inherited by tumor cells from their cell of origin have clinical relevance for many cancer types^2^.

B-cell lymphomas, a heterogeneous set of malignancies arising from different stages of B-cell maturation in lymph nodes, have been well studied in this regard. In the B-cell maturation process, naïve B cells enter the germinal center (GC) upon T-cell dependent activation, where they proliferate and undergo somatic hypermutation in the dark zone (DZ) before migrating to the light zone (LZ) for antigen presentation by T follicular helper cells (TFH) and follicular dendritic cells (FDC), leading to differentiation into memory B cells (Mem) or plasma cells^3,4^. B-cell lymphoma subtypes include typically pre-GC mantle cell lymphoma (MCL), GC diffuse large B-cell lymphoma (DLBCL, GCB) and follicular lymphoma (FL), post-GC or activated B-cell DLBCL (DLBCL, non-GCB/ABC), as well as memory-origin marginal zone lymphoma (MZL)^4–7^. These B-cell lymphoma subtypes have variable clinical courses, ranging from the more indolent FL and MZL to the more aggressive MCL and DLBCL^8^. Following standard treatment with chemoimmunotherapy, ABC-like-DLBCL shows more frequent relapse compared to GCB-like-DLBCL. This association between DLBCL tumor maturation state and clinical outcomes has more recently been extended beyond the ABC vs GCB dichotomy to across the maturation spectrum^9^.

In addition to the cell of origin, cancer pathogenesis and treatment response are also shaped by intratumor heterogeneity^13^. Intratumor heterogeneity can manifest at genetic, epigenetic, cellular phenotype, and microenvironment levels, contributing to complex tumor architecture, evolution, and adaptive resistance mechanisms^10,14^. Greater intratumor heterogeneity has been associated with a worse prognosis, due to the increased likelihood of therapy-resistant subclones^15–17^. In B-cell lymphoma, substantial heterogeneity has been observed in the tumor microenvironment, including in stromal^18^ and T-cell infiltration patterns^19^, while intratumor transcriptional subpopulations have shown differences in their treatment sensitivities^10^.

Understanding the drivers of intratumor heterogeneity promises to improve our understanding of cancer pathogenesis, evolution, and resistance mechanisms. While cell of origin significantly contributes to *inter*tumor heterogeneity, the role of differentiation processes like B-cell maturation in *intra*tumor heterogeneity is unclear. Dedifferentiation into cancer stem cells and transdifferentiation across lineages have been recognized in several other cancer types^20,21^, and limited plasticity beyond the cell of origin has been noted in 4 FL^22^ and 4 DLBCL^23^ tumors. Nevertheless, the prevailing paradigm in cancer pathogenesis remains that cancer cells avert normal cellular differentiation to avoid terminal differentiation and maintain proliferation^24–26^. Through transcriptional, proteomic, epigenetic, genetic, and spatial profiling of MCL, FL, DLBCL, MZL, and reactive lymph nodes (rLN) across 51 patients, representing the largest cross-entity single-cell atlas of B-cell lymphoma tumors to date, we show that cellular differentiation continues in malignancy, driving intratumor heterogeneity and response.

## Methods

### Study outline

This study was approved by the University of Heidelberg’s Ethics Committee (S-254/2016), with informed consent obtained from all participants. Lymph node (LN) samples from 51 patients (8 MCL; 12 FL; 5 GCB DLBCL; 7 ABC DLBCL; 11 MZL; 8 rLN, Supplemental Table 1) were processed and cryopreserved using published protocols^27^. Cellular Indexing of Transcriptomes and Epitopes by Sequencing (CITE-Seq)^28^ and Chromium Single Cell Immune Profiling were performed as per the manufacturer’s (10X Genomics) instructions and analyzed with the *Seurat*^29^ R package. Highly multiplexed histochemistry (CODEX) was performed as previously described^30^. Antibody panels are outlined in Supplemental Tables 2-4. Transcription factor activity and copy number variants were inferred with the *SCENIC*^31^ and *copykat*^32^ packages. Targeted DNA sequencing was performed on the targets outlined in Supplemental Table 7. Gene set enrichment analysis was performed with the *fgsea*^33^ R package using the MSigDB hallmark gene set collection^34^. For survival analysis, the maturation state composition of 507 FL and DLBCL tumors was deconvoluted from the Loeffler-Wirth et al. 2019^35^ microarray dataset using CIBERSORTx^36^. Cox proportional hazards models and Kaplan-Meier curves were generated with the *survival*^37^ and *survminer*^38^ R packages. See the Extended Methods section of the Supplemental Data for a more detailed description of the methods.

### Data availability

All single-cell sequencing data has been deposited at the European Genome-phenome Archive (EGA), which is hosted by the EBI and the CRG, under accession number EGAS50000000342. Further information about EGA can be found at https://ega-archive.org and “The European Genome-phenome Archive of human data consented for biomedical research”. Highly multiplexed immunofluorescence images are available in the [EMBL-EBI BioImage Archive](https://www.ebi.ac.uk/bioimage-archive/) under accession ID S-BIAD1090.

### Code availability

Code scripts used for data processing, analyses, and figure generation are publicly available at https://github.com/dnfitzgerald/BNHL-atlas.

## Results

### A single-cell B-cell maturation reference map in reactive lymph nodes

To characterize nodal B-cell maturation states in the non-malignant context, we performed Louvain clustering on the rLN single-cell transcriptomic data, obtaining 16 clusters that we assigned to naïve, germinal dark zone centroblasts (DZ), light zone centrocytes (LZ), IgD+/IgM+ memory (Mem IgM), class-switched memory of predominantly IgG class (Mem IgG), and plasma maturation states based on the presence of established markers^3,4,9,39–43^(Supplemental Table 5, Fig. 1b-c). These B-cell maturation state annotations were validated with FACS sorting and RNA sequencing, which confirmed the presence and gene expression profiles of the identified states (Supplemental Fig. 1, Fig. 1d). We observed that the rLN samples were composed of 60% memory (split equally between IgD+/IgM+ (Mem IgM), and class-switched (Mem IgG)), 30% naïve, 6% light zone centrocytes (LZ), 3% dark zone centroblasts (DZ), and 2% plasma cells.

**Fig. 1:**
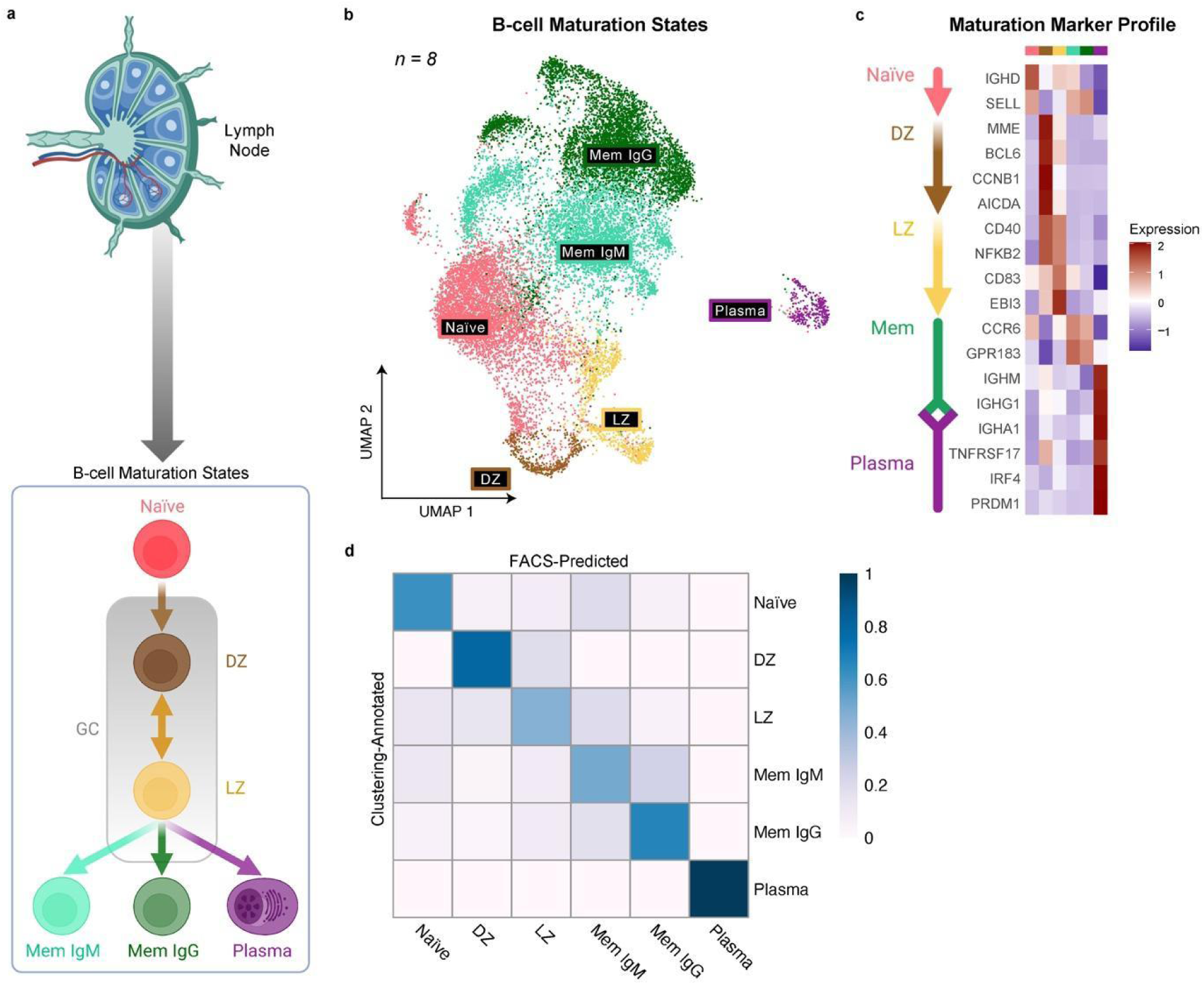
A single-cell B-cell maturation reference map in reactive lymph nodes. **a**, Schematic of the B-cell maturation trajectory in the lymph node. The labeled populations represent the B-cell maturation states characterized by FACS and CITE-Seq. Illustrations were created with BioRender.com^69^. **b**, Transcriptomic UMAP of the rLN reference CITE-Seq dataset (8 samples) labeled by the B-cell maturation states in a. Transcriptomic clusters were assigned to maturation states based on their expression of the maturation markers in Supplemental Table 5. **c**, Heatmap showing the z-scored average expression of a subset of markers for each maturation state annotated in the reference CITE-Seq dataset. **d**, Confusion matrix of cells’ maturation state labels annotated by maturation marker profiling of transcriptomic clusters (y-axis) and predicted by a logistic regression^70^ classifier trained on RNA-sequencing data from B-cell maturation states sorted with FACS (x-axis) (Supplemental Fig. 1). The color scale shows probability estimates for each class. Maturation state annotations: Naïve = Naïve B cells, DZ = Centroblasts from the dark zone of the germinal center, LZ = Centrocytes from the light zone of the germinal center, Mem IgM = IgD+ and IgM+ memory B cells, Mem IgG = class-switched (IgG+ or IgA+) memory B cells, Plasma = plasma cells.

### Divergence of B-cell maturation in tumors

Having characterized B-cell maturation states in non-malignant lymph nodes, we profiled these states in the 43 B-cell lymphoma samples (8 MCL, 12 FL, 5 GCB DLBCL, 7 non-GCB DLBCL, and 11 MZL). We identified malignant cells based on light chain restriction^44^, with transcriptional clusters having >75% kappa or lambda light chains considered malignant. BCR profiling in 8 samples confirmed each tumor had a single expanded B-cell receptor clone with a restricted light chain. Non-malignant B cells, mainly naïve B cells, comprised a median of 6% [0%-94%] of all B cells across tumor samples (Supplemental Fig. 2).

To classify B-cell maturation states in tumors, we used mutual nearest neighbors and canonical correlation analysis^45^ to map maturation states from the rLN reference dataset to each tumor. The classified maturation states in malignant cells matched their maturation marker profiles. We also calculated maturation state gene signature scores using the 50 most differentially expressed genes from a published tonsil dataset^9^, and these scores in tumors reflected those in rLN samples (Supplemental Fig. 3a-b).

Hereon focusing on malignant B cells, we found that tumors displayed a spectrum of maturation states, indicating plasticity and divergence from a common cell of origin (Fig. 2a-b, Supplemental Fig. 4). This indicates that B-cell maturation remains plastic and divergent in malignancy, rather than fixed.

**Fig. 2:**
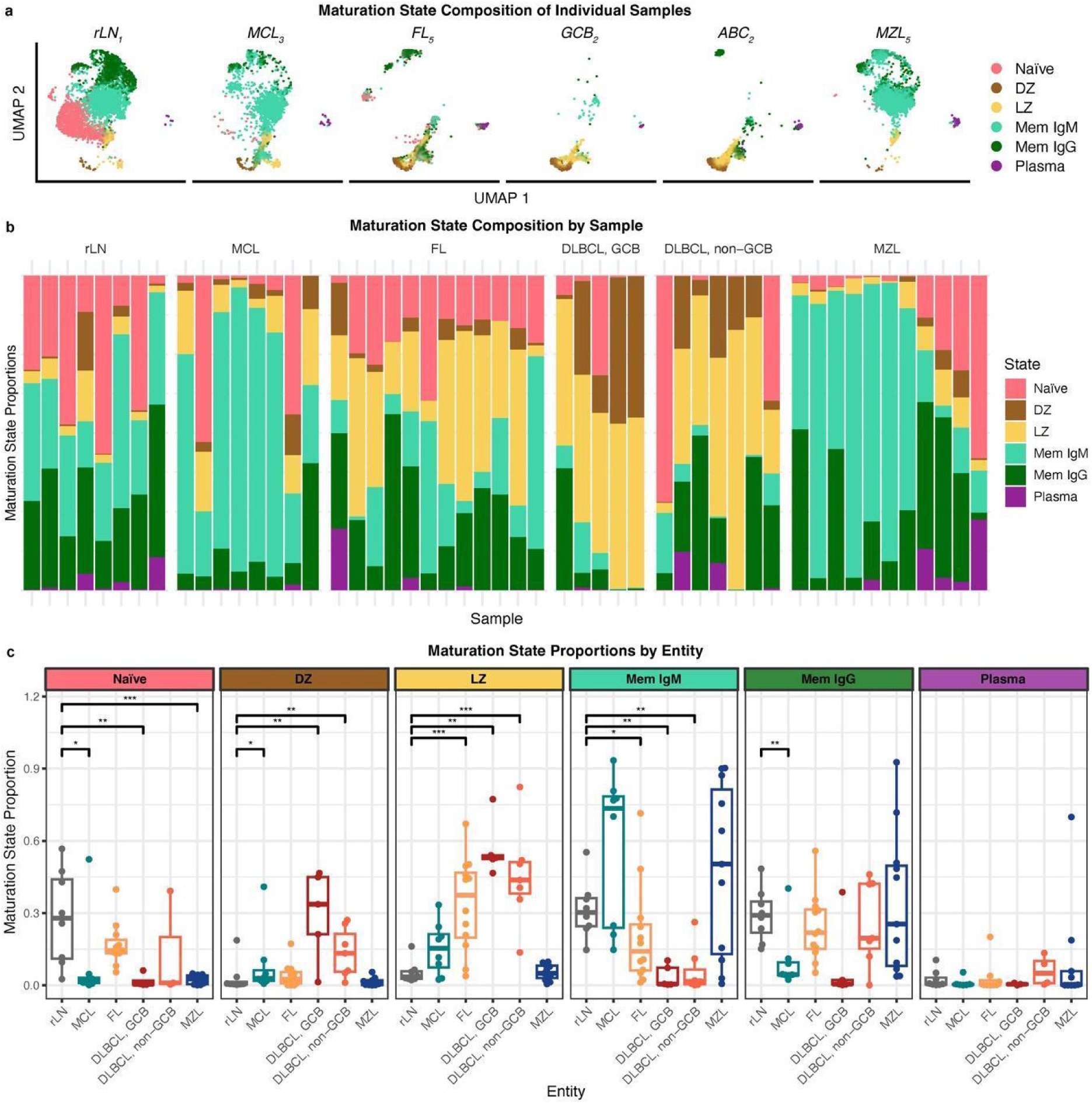
Tumors consist of multiple B-cell maturation states. **a**, Transcriptomic UMAPs for individual samples from each subtype, labeled by B-cell maturation states assigned by label transfer from the reactive lymph node reference (Fig. 1b). Only malignant cells are shown for tumor samples. **b**, Maturation state composition of all samples (n=51) split by subtype and ordered by days since diagnosis. **c**, Box plot of the proportion of each maturation state in each subtype. Each data point is a sample, grouped by subtype. The Wilcoxon signed-rank test^71^ (two-sided) was performed for each maturation state’s proportion between reactive lymph nodes and each subtype: p<0.05 (*), p<0.01 (**), p<0.001 (***). Centerline = median, box limits = upper and lower quartiles, whiskers = 1.5x interquartile range. rLN = reactive lymph nodes, MCL = mantle cell lymphoma, FL = follicular lymphoma, DLBCL = diffuse large B-cell lymphoma (GCB = germinal center, non-GCB/ABC = activated B cell), MZL = marginal zone lymphoma. See Fig. 1 for maturation state annotations.

### Maturation states are a source of intratumor and intertumor heterogeneity

We observed a characteristic spectrum of maturation states in each subtype; the predominant states reflected their associated cell of origin, such as GC states in DLBCL GCB and memory states in MZL, and co-existed with states both upstream and downstream in the maturation process. We also observed substantial variation in maturation state proportions between samples of the same subtype (Fig. 2b). B-cell maturation states contributed significantly to both intratumor and intertumor heterogeneity, with subtype prediction accuracy reflecting this complexity (Supplemental Fig. 3c).

FL and non-GCB DLBCL tumors showed diverse mixtures of GC (DZ and LZ) and post-GC (memory and plasma) states, suggesting that, like DLBCL, FL may also be capable of transformation into post-GC phenotypes. Both DLBCL and FL showed significant enrichment in the LZ state compared to rLN controls (p < 0.01, Wilcoxon signed rank test), but FL tumors were not enriched for the DZ state (Fig. 2b), implying that DLBCL better maintains a DZ phenotype. Continuation of somatic hypermutation and proliferation in the DZ state may promote DLBCL’s characteristically more aggressive disease course^46^.

We explored phenotypic diversity in B-cell lymphoma tumor cells by conducting unsupervised multimodal subpopulation mapping on the full CITE-Seq B-cell dataset (51 samples, 154,282 cells). Using Multi-Omic Factor Analysis (MOFA)^47^, we identified 26 multimodal subpopulations, mostly segregating by maturation state, underscoring differentiation as a major driver of tumor variation (Supplemental Fig. 5a-e). The greatest heterogeneity was observed among memory B-cell states, with subpopulations segregating by B-cell lymphoma subtype. Features of these subpopulations pointed to known pathogenic mechanisms, such as CCND1 overexpression in MCL^48^ (Supplemental Fig. 5f-g).

### Cell-of-origin classification reveals multiple subtypes within each tumor

Next, we examined the implications of intratumor variation in maturation states for established diagnostic cell-of-origin (COO) classifiers. Applying the Lymph2Cx gene expression classifier^49^ to single-cell RNA-sequencing (scRNA-seq) data from DLBCL tumors revealed that most tumors consisted of multiple COO subtypes (GCB, unclassified, and ABC), rather than a single GCB or ABC class per tumor. This indicates a more complex pathology in DLBCL, with a mixture of COO subtypes within each tumor (Fig. 3a-b). This observation was confirmed using the Tally DLBCL COO classifier, which employs immunohistochemistry markers (e.g., CD10, GCET1, LMO2 for GCB; MUM1, FOXP1 for non-GCB) to classify tumors^50^. Both classifiers showed that GCB DLBCL often contains ABC-like cells, consistent with a portion of GCB-origin tumor cells transitioning to an ABC subtype (Fig. 3d-e).

**Fig. 3:**
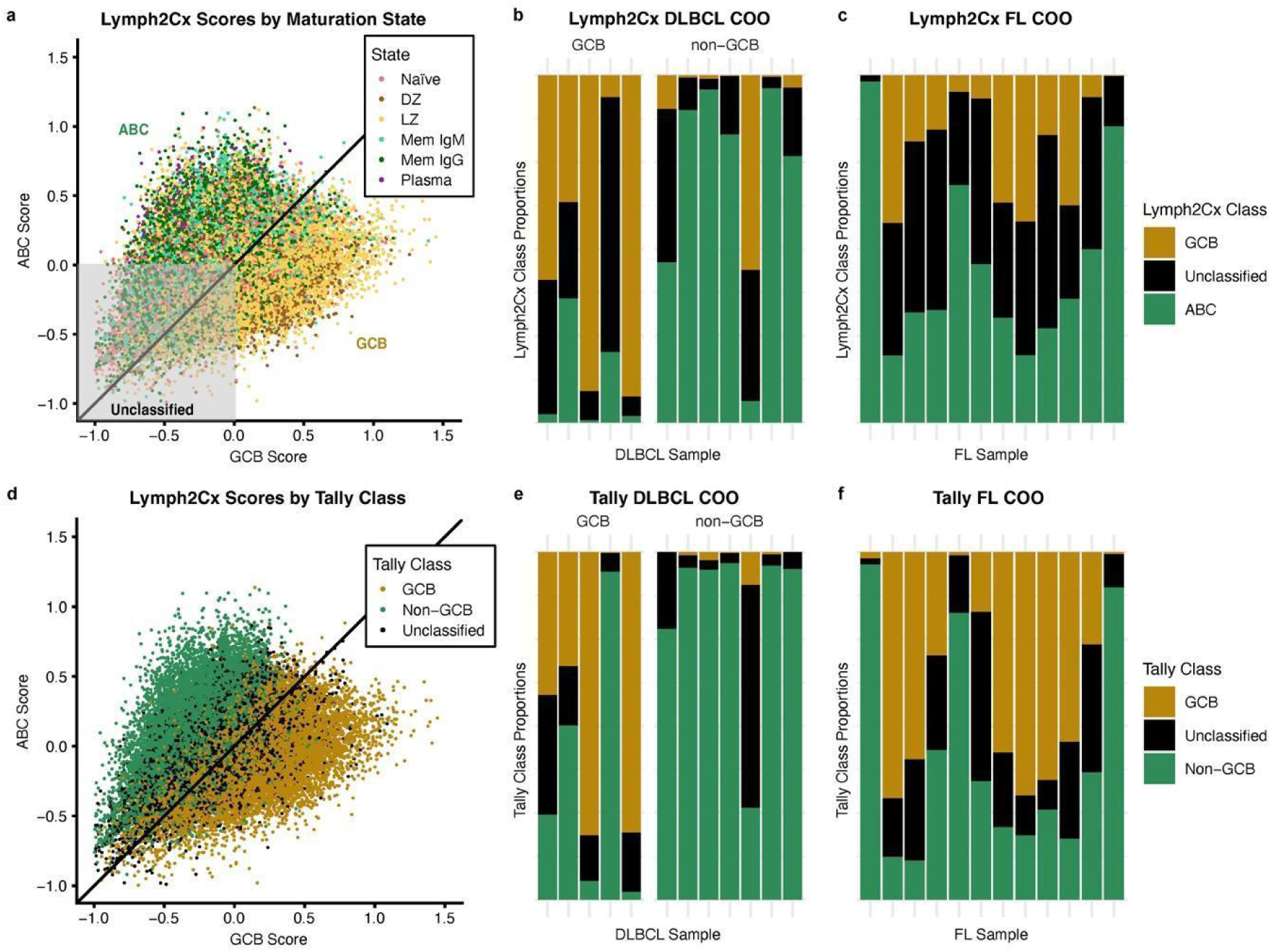
Cell-of-origin classification reveals multiple subtypes within each tumor. **a**, Normalized GCB and ABC scores for each DLBCL and FL tumor cell determined by the Lymph2Cx cell-of-origin classifier, labeled by B-cell maturation state. The diagonal line divides cells by their classification (above = ABC, below = GCB). Cells without ABC or GCB gene set average expression above the housekeeping genes were unclassified (grey box). **b-c**, Lymph2Cx class proportions among the tumor cells of DLBCL samples faceted by GCB or non-GCB diagnosis (b) and FL samples (c). **d**, Normalized GCB and ABC scores for each DLBCL and FL tumor cell determined by the Lymph2Cx classifier, as shown in (a), labeled by Tally class. **e-f**, Tally class proportions among the tumor cells of DLBCL samples faceted by GCB or non-GCB diagnosis (e) and FL samples (f). FL = follicular lymphoma, DLBCL = diffuse large B-cell lymphoma (GCB = germinal center, non-GCB = non-germinal center), COO = cell of origin. See Fig.1 for maturation state annotations.

Interestingly, FL tumors, typically not associated with an ABC cell of origin, displayed both GCB and ABC classes at the single-cell level in 10 out of 12 tumors when using Lymph2Cx and Tally classifiers. This suggests the emergence of an ABC-like population within FL tumors (Fig. 3c+f).

### Tumor maturation state composition changes over time

We hypothesized that intratumor heterogeneity introduced by B-cell maturation might drive tumor evolution, with shifts in maturation state composition over time due to disease progression or treatment selection. This hypothesis was supported by observed increases in class-switched memory (Mem IgG) and plasma states in MZL and an increase in the ABC COO subtype in FL samples collected later in the disease course (Fig. 2b). In longitudinal lymph node samples from three B-cell lymphoma patients, we observed shifts in maturation state composition, such as a decrease in plasma cells and an increase in Mem IgG in an MZL tumor that relapsed 15 months after 6 cycles of obinutuzumab and bendamustine (Supplemental Fig. 6a). Similarly, plasma cells increased after 11 months in a GCB DLBCL tumor that relapsed following CAR-T therapy (axicabtagene ciloleucel) (Supplemental Fig. 6b). However, little change was observed after 2 months in another DLBCL patient who relapsed to the same CAR-T therapy, suggesting that these effects may be time-dependent (Supplemental Fig. 6c).

### Transcription factor signatures of maturation states maintain differential activity in malignancy

Transcription factors (TFs) are key mediators of the B-cell maturation process, although alteration of the epigenomic landscape has been described in the pathogenesis of B-cell lymphoma^5,39,51–53^. We investigated whether maturation-associated TFs were maintained in malignancy, which could enable a tumor to diverge into multiple maturation states despite epigenomic dysregulation. Using the *SCENIC* workflow^31^ on scRNA-seq data (51 samples), we identified coexpression modules between TFs and target genes, scoring TF activity based on module expression. Signature TFs were defined by differentially expressed target genes between maturation states in rLN samples (BH-adjusted p-value < 10e-16, average log fold-change > 0.4). These maturation TF signatures were largely conserved across B-cell lymphoma subtypes, though this conservation was partially lost in the LZ state of DLBCL, where TFs like *IRF8, NFKB1, HIVEP3*, and *REL*, elevated in rLN LZ, showed higher activity in the memory state (Fig. 4a). Variation in memory signature TF activity across subtypes (e.g., *IRF7, STAT1, IRF9*) corresponded with the diversity of nodal memory B-cells, which are less dependent on maturation GRNs after completing the maturation process. TFs with the most significant activity differences in malignancy were linked to B-cell development, activation, and differentiation, including *KLF3* in naïve and memory B cells, *MAZ* and *HDAC2* in GC B cells, *TBL1XR1* in memory, and *XBP1* in plasma cells (Fig. 4b, Supplemental Fig. 7).

**Fig. 4:**
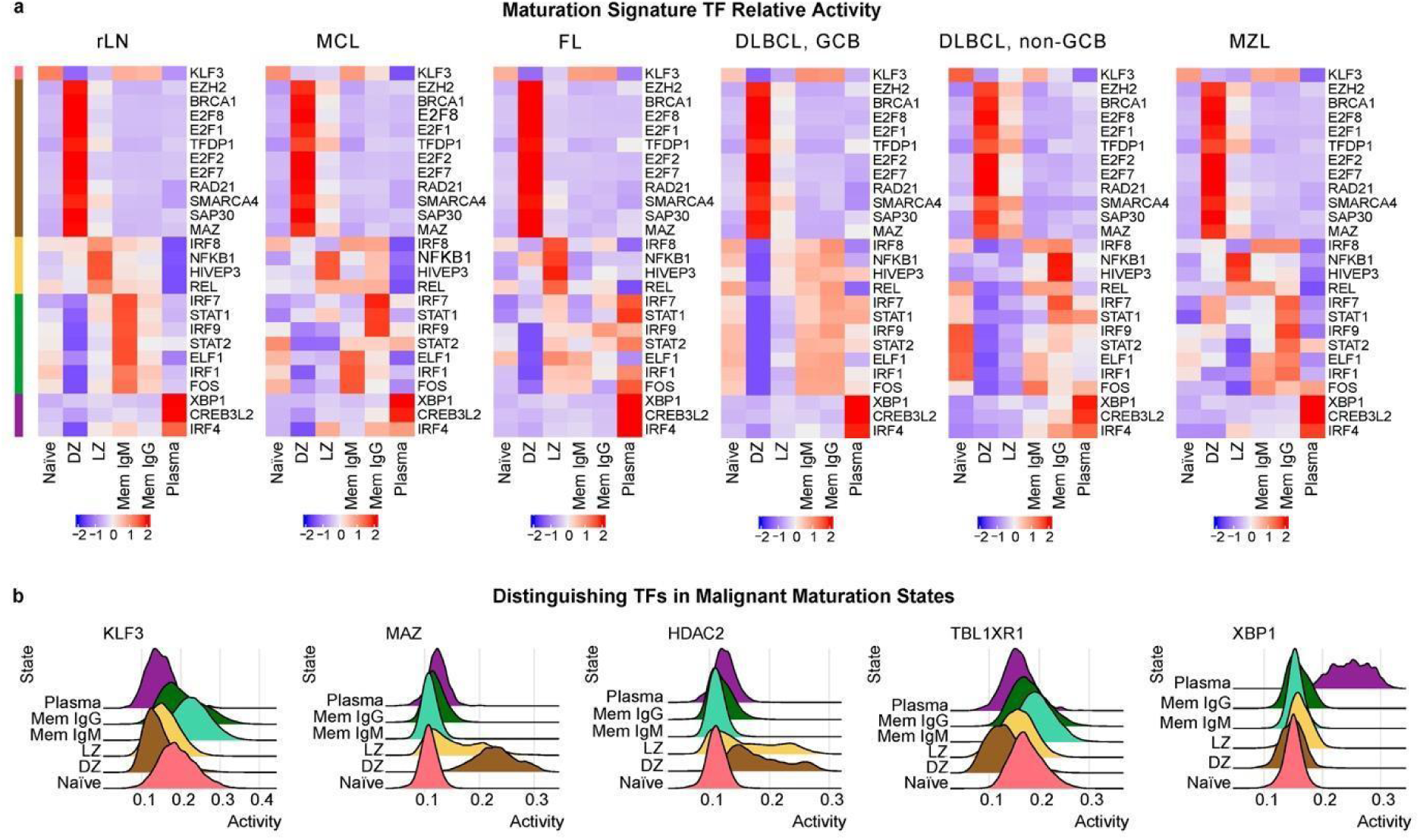
Maturation signature TFs maintain differential activity in malignancy. **a**, Scaled activity of transcription factors (TFs) across B-cell maturation states in each subtype inferred using the *scenic* python package^31^; only TFs significantly enriched (log2 fold-change >0.4, p < 10e-16 as determined with the *MAST* R package^72^) for B-cell maturation stages target genes in reactive lymph node samples are shown. TFs (y-axis) are ordered by the maturation state in reactive lymph nodes in which they are enriched. Non-malignant cells in tumor samples were excluded based on light-chain restriction. **b**, Density plots comparing the activity distribution of a subset of distinguishing transcription factors in each tumor maturation state in malignant cells aggregated from all tumor samples (n = 43). rLN = reactive lymph node, MCL = mantle cell lymphoma, FL = follicular lymphoma, DLBCL = diffuse large B-cell lymphoma (GCB = germinal center, non-GCB = non-germinal center), MZL = marginal zone lymphoma. See Fig. 1 for maturation state annotations.

### Maturation states occupy distinct tumor microenvironments

The follicular architecture of lymph nodes is central to the B-cell maturation process. Therefore, we sought to understand the spatial context of intratumor maturation states and whether spatial niches could still facilitate maturation in malignancy. We characterized the spatial distribution of cell types with CODEX^30^ (52 features, Supplemental Table 7) for 19 samples (29 slides) in the CITE-Seq cohort. Immune cell types were defined according to previously published nodal cell type annotations^19^ (Supplemental Fig. 9a). B-cell maturation state labels were transferred from the CITE-Seq to the CODEX dataset via shared protein features (n=28), yielding a high correlation in maturation state proportions between the datasets across samples (median R=0.91, p=0.011) (Supplemental Fig. 8). Tumor microenvironment composition varied significantly across tumors and subtypes, with MCL and DLBCL typically losing normal follicular structure. However, tumor cells of different maturation states still tended to segregate spatially (Supplemental Fig. 9b). For instance, FL1 showed expanded follicles with DZ and LZ states surrounded by memory and plasma cells. At the same time, MCL1 exhibited spatially distinct LZ and memory states, but without typical follicular structure. ABC2 displayed a spread of several maturation states with little spatial distinction, consistent with DLBCL’s diffuse nature (Fig. 5a).

**Fig. 5:**
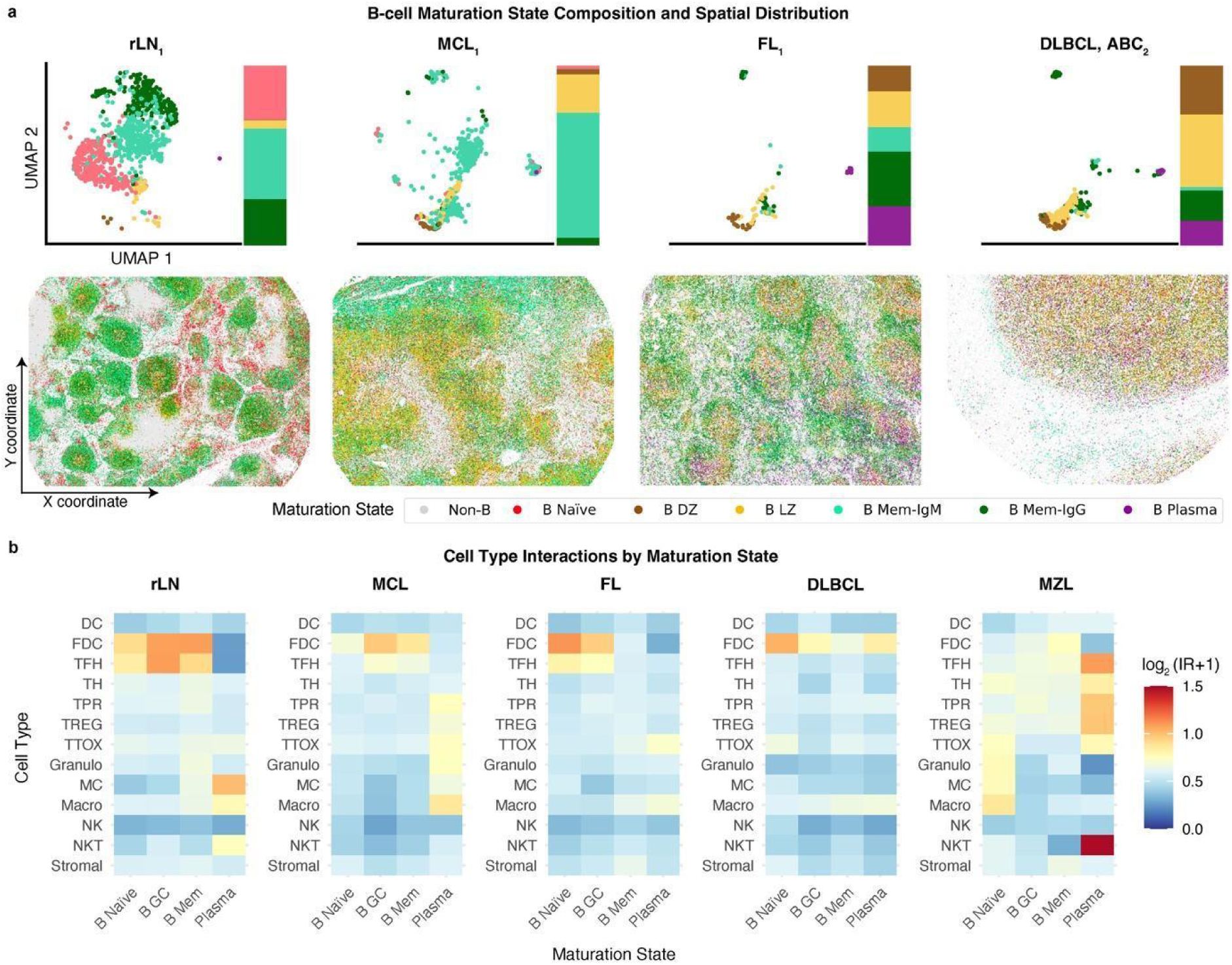
Maturation states occupy distinct tumor microenvironments. **a**, UMAP of scRNA-seq data labeled by B-cell maturation states mapped from the reactive lymph node reference with B-cell maturation state proportions (top), and spatial distribution of B-cell maturation states in a reactive lymph node (rLN), follicular (FL), mantle cell (MCL), and non-germinal center diffuse large B-cell lymphoma (DLBCL, non-GCB) sample from CODEX^30^ images (52 markers) on FFPE tissue sections (bottom). See Supplemental Fig. 9 for the distribution of maturation states and cell types on all CODEX slides (n=29). **b**, Log_2_ of the pairwise cell-cell observed over expected interaction ratios (IR) between B-cell maturation states and all other cell types in each subtype, with a pseudocount of 1. Spatial interactions were determined with Delaunay triangulation. A higher ratio is associated with increased proximity. B Naïve = naïve B cells, B DZ = centroblasts from the dark zone of the germinal center, B LZ = centrocytes from the light zone of the germinal center, B Mem IgM = IgD+ and IgM+ memory B cells, B Mem IgG = class-switched (IgG+ or IgA+) memory B cells, B Plasma = plasma cells, CD4T_naive = naive CD4+ T-cells, CD8T_naive = naive CD8+ T-cells, TH_memory = memory helper T-cells, TTOX_memory = memory cytotoxic T-cells, TTOX_exh = exhausted cytotoxic T-cells, NKT = natural killer T-cells, TFH = follicular helper T-cells, TPR = proliferating T-cells, TREG = regulatory T-cells, FDC = follicular dendritic cells, DC = dendritic cells, Macro = macrophages, Stromal = stromal cells, NK = natural killer cells, MC = monocytes, Granulo = granulocytes.

We explored spatial niches using K nearest neighbor (KNN, K = 20) graph-based cellular neighborhood (CN) analysis, identifying 11 CNs defined by enriched cell types (Supplemental Fig. 10). CNs common in rLNs were also present in tumors, including T-cell zones and a stromal zone bordering B cells. However, most malignant B cells were found in CNs distinct from the follicular CNs seen in rLN samples. Each intratumor maturation state was dominant in a different CN and associated with different immune compositions, such as cytotoxic T-cells and macrophages in plasma zones (Supplemental Fig. 10).

Heterogeneous microenvironments may promote maturation state divergence by enabling or restricting specific cellular interactions critical to maturation. FDCs and TFH cells were enriched in GC-predominant tumor CNs, potentially supporting the ongoing differentiation of malignant GC cells into post-GC memory or plasma cells (Supplemental Fig. 10b). Pairwise cell-cell interaction ratios (IR) between B-cell maturation states and other cell types indicated that GC B-cell interactions with FDC and TFH were partially maintained in tumors, allowing the GC reaction to continue in malignancy. This was less evident in the more diffuse DLBCL and generally post-GC MZL, possibly explaining their lower diversity of maturation states compared to FL (Fig. 2b).

### Genetic variants among intratumor maturation states

Several characteristic features of B-cell maturation states are known to be commonly aberrated in B-cell lymphoma (eg. *BCL6, IRF4,* and *PRDM1*)^3,54^. We considered whether intratumor maturation states may be associated with subclonal genetic variation. We profiled genetic variants, including single-nucleotide variants (SNVs), insertions and deletions, and copy number variants (CNVs) from targeted DNA-sequencing data in bulk tumor samples. We inferred CNVs at the single-cell level from gene expression data using the *copykat* method^32^. Distinct CNVs were observed within individual tumors, often confined to specific maturation states. For example, in FL7, most GC tumor cells displayed aneuploidy compared to predominantly diploid non-GC cells (Fig. 6a-d). This tumor exhibited a 6p22.2 copy number gain, associated with FL progression^55^, restricted to DZ tumor cells (Fig. 6c), suggesting that this gain may either lock cells in the DZ stage or that DZ cells are more prone to acquiring such variants.

**Fig. 6:**
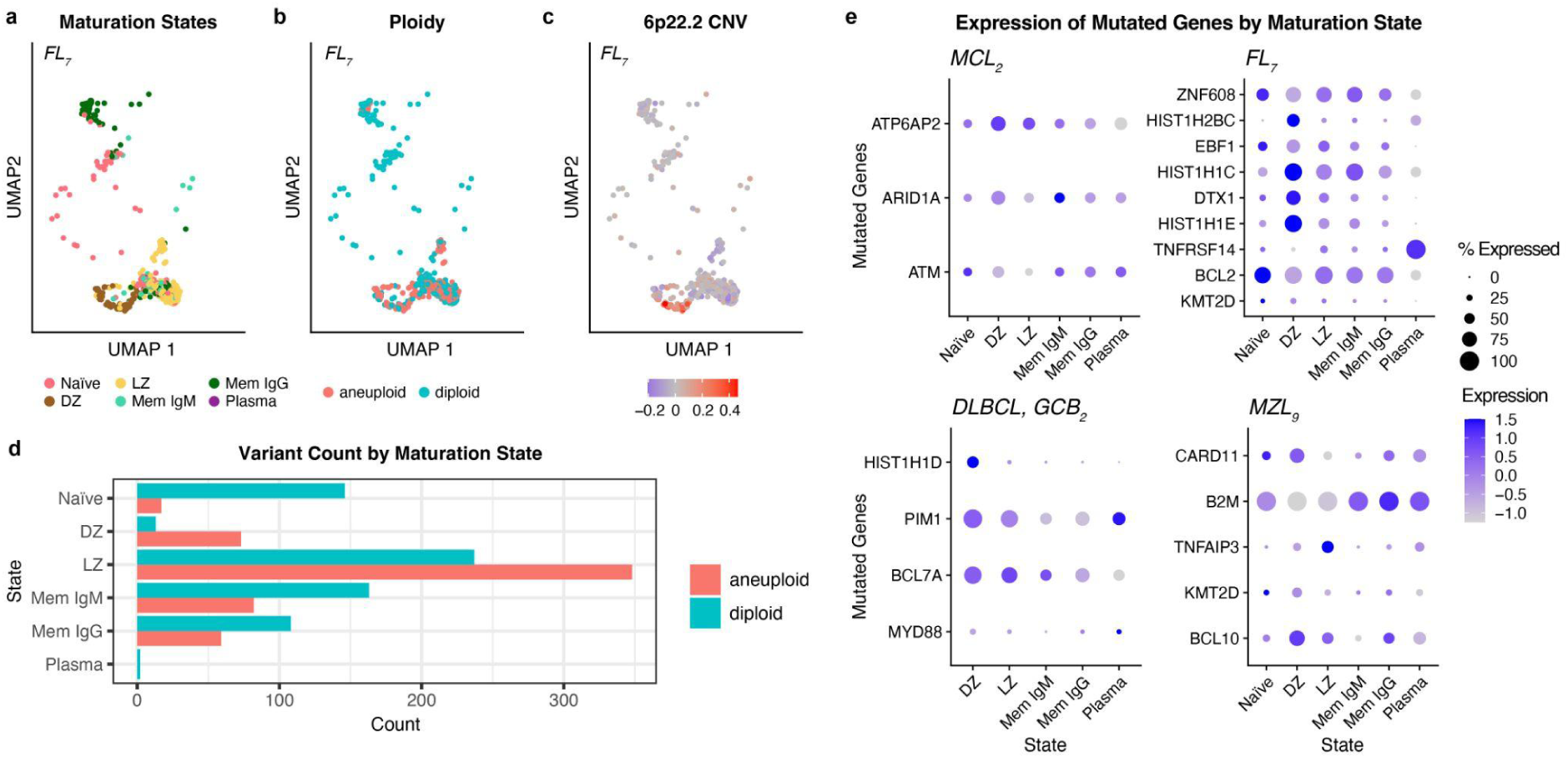
Genetic variants among intratumor maturation states. **a-c**, Reference-based UMAP of malignant cells from FL7 labeled by (a) B-cell maturation state, (b) diploid or aneuploid status, and (c) copy number variation (CNV) in chromosomal position 6-26329011 (cytogenetic band 6p22.2) among tumor cells. CNV was inferred from gene expression with the *copykat* R package^32^. **d**, Frequencies of the aneuploid and diploid variants in each intratumor maturation state in FL7. **e**, The average expression (z-scaled) and percentage of cells expressing genes with non-silent mutations detected with targeted DNA sequencing (Supplemental Fig.11) in samples from mantle cell lymphoma (MCL), follicular lymphoma (FL), diffuse large B-cell lymphoma (DLBCL) and marginal zone lymphoma (MZL). See Fig. 1 for maturation state annotations.

Genes harboring non-silent mutations also differed in expression between intratumor maturation states. For instance, an MCL tumor with multi-hit *ATM* mutations showed restored expression in post-GC states, while an FL tumor with *HIST1H1E* and *DTX1* mutations showed reduced expression in post-DZ states (Fig. 6e). These findings are consistent with a recent study in which we observed differential mutation abundance between maturation states in the same tumor (eg. BCL2 variants enriched in LZ vs DZ)^56^.

### Tumor maturation state composition predicts survival

We investigated the impact of intratumor B-cell maturation state heterogeneity on survival by deconvoluting maturation states with CIBERSORTx^36^ from a microarray dataset containing 430 DLBCL, 145 FL, and 48 mixed FL/DLBCL tumors that were heterogeneously treated pre-Rituximab era^35^. Cox proportional hazards analysis revealed that IgM memory and DZ states were associated with poorer survival, while LZ and naïve states were linked to prolonged survival (Fig. 7a). Gene set enrichment analysis showed that IgM memory and DZ states exhibited increased proliferation and cell growth (Myc, E2F, and mTOR) signaling and DNA repair, indicative of a more aggressive and resistant phenotype^57,58^ (Fig. 7b-c). The DZ state showed reduced dependence on canonical TNF, JAK-STAT, and KRAS signaling compared to other tumor populations, which could enable them to escape the typical immune response and therapeutic interventions (Fig. 7b-c). IgM memory B cells can class switch to IgG/IgA memory, differentiate into plasma cells, or re-enter the germinal center reaction to undergo further somatic hypermutation and proliferation^59^. This special plasticity could give IgM memory tumor cells an evolutionary advantage through adaptation, thus driving poorer survival outcomes.

**Fig. 7:**
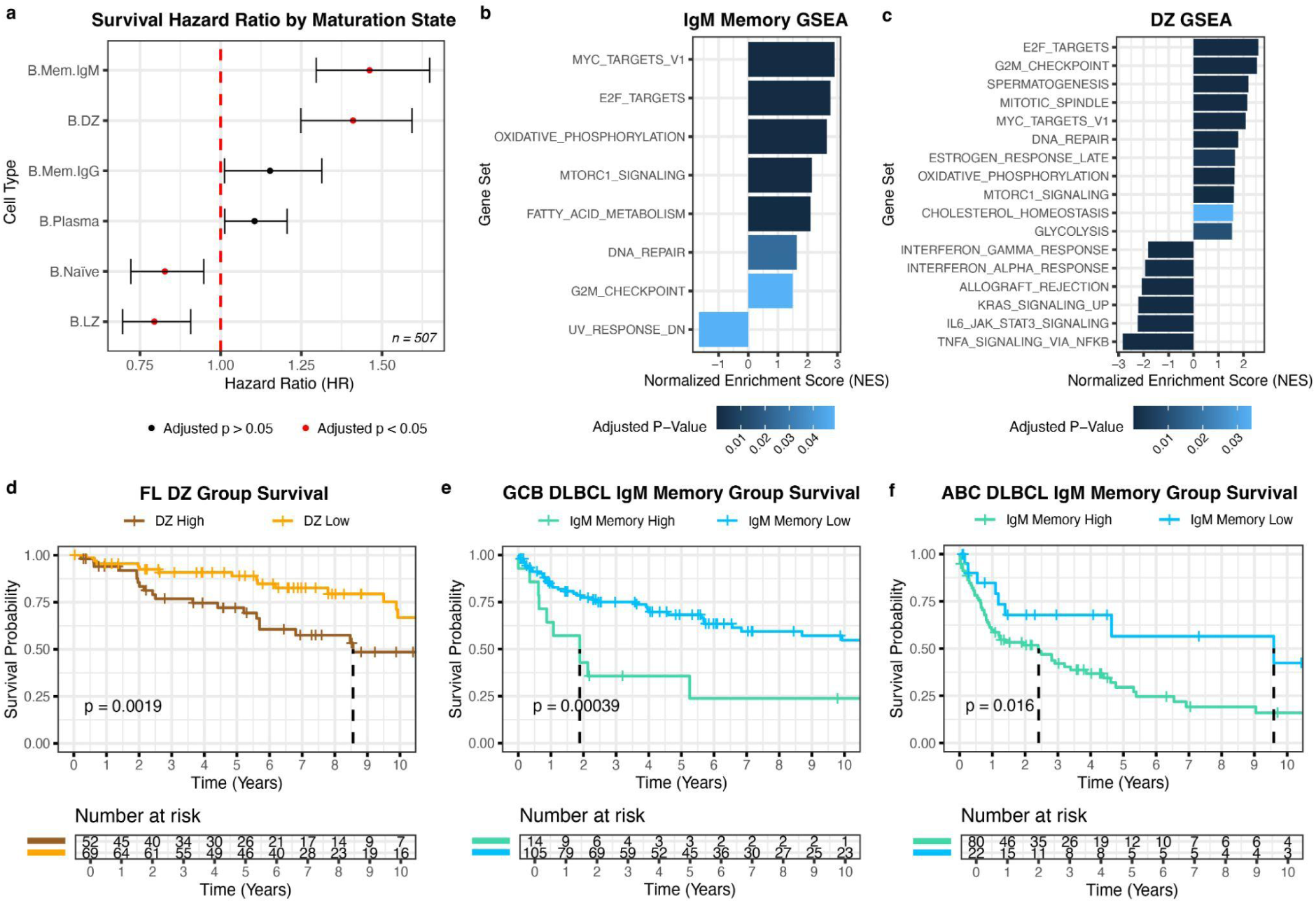
Tumor maturation state composition predicts survival. **a**, Hazard ratios for each maturation state (for 1 standard deviation) across tumors and their 95% confidence intervals are shown with respect to overall survival from 507 tumors (119 DLBCL GCB, 102 DLBCL ABC, 79 unclassified DLBCL, 44 double-hit DLBCL, 94 BCL2-break positive FL, 25 BCL2-break negative FL, 2 BCL-break unknown FL, and 42 mixed FL/DLBCL (FL grade 3B)). Significant hazard ratios (adjusted p<0.05, Log-Rank test, Benjamini-Hochberg adjusted^73^) are shown in red. See Fig. 1 for B-cell maturation state annotations. **d-f**, Kaplan-Meier curves are shown for **d**, DZ high and low (>/< 1.9%) subgroups of FL, **e**, IgM memory high and low (>/< 2.66%) subgroups of DLBCL GCB, **f**, IgM memory high and low (>/< 1.4%) subgroups of DLBCL ABC. P-values were determined with the log-rank test and 95% confidence intervals are shown.

We identified distinct survival groups based on maturation state proportions. In FL, tumors enriched for DZ cells showed a more aggressive course (median survival = 8.6 vs. 13 years; χ² = 9.6, df = 1, p = 0.0019). In DLBCL, IgM memory high and low subgroups had striking survival differences in both GCB and ABC subtypes (GCB: median = 1.9 vs. >10 years, χ² = 12.6, df = 1, p = 0.00039; ABC: median = 2.4 vs. 9.6 years, χ² = 5.8, df = 1, p = 0.016). These results suggest that intratumor maturation state profiling can identify high-risk patient groups within B-cell lymphoma subtypes.

## Discussion

The prevailing view in cancer pathogenesis is that normal cellular differentiation is averted in malignancy, while cancer types and subtypes are tied to distinct cell types and states^1,4,5,24,54^. In this paradigm, B-cell lymphoma subtypes are associated with different stages of the B-cell maturation lineage^4,5,54^. In contrast, our study reveals that ongoing differentiation drives intratumor heterogeneity in MCL, FL, DLBCL, and MZL, with tumors exhibiting a spectrum of B-cell maturation states.

We show that ongoing B-cell maturation is a major driver of intratumor heterogeneity across B-cell lymphoma subtypes. Intratumor heterogeneity in maturation states transcends subtype boundaries, which are tied to different cell types of origin. This deviation in maturation is bidirectional; post-GC states are observed in GC tumors (eg., FL), and GC states are observed in post-GC tumors (eg., DLBCL non-GCB and MZL). This raises two potential roles of maturation in B-cell lymphoma pathogenesis; it may either enable additional tumor maturation states to evolve from the subtype-determining cell of origin or drive the emergence of the predominant subtype-associated tumor maturation state from a cell of origin earlier in the maturation process.

As B-cell lymphoma subtypes^8^ associated with different maturation states (e.g. GCB vs ABC in DLBCL)^9,60^ have distinct clinical outcomes, variation in tumor maturation state composition may explain the variable disease courses among patients. This is especially plausible given that the Lymph2Cx and Tally DLBCL cell-of-origin subtype classifiers, which distinguish GCB and ABC DLBCL subtypes with distinct prognoses, reveal the presence of both GCB and ABC classes within individual tumors, even in FL where a post-GC subtype has not yet been described. Thus, intratumor heterogeneity in maturation may require revising B-cell lymphoma subtypes to account for mutability and multiplicity of maturation states within tumors. The identification of survival-based subgroups of FL, GCB DLBCL and ABC DLBCL based on the abundance of DZ and IgM memory states suggests that tumor maturation state profiling could be used for patient stratification and management. Future studies may link this intratumor heterogeneity to evolving molecular classifications of B-cell lymphomas (e.g., DZsig^61^ and LymphGen^62^) and study cancer cell type diversity and its consequences across cancers.

The plasticity in B-cell maturation observed in malignancy would enable shifts in intratumor maturation states over time, promoting tumor evolution and expanding the selection pool for treatment resistance. Understanding the link between intratumor maturation states and resistance mechanisms could inform treatment stratification strategies. Targeting vulnerabilities in specific maturation states, or blocking differentiation to more resistant states, could offer therapeutic opportunities. Some pathways and gene regulatory networks (e.g., XBP1, HDAC2^63^) already have known ligands or approved drugs, though care is needed to avoid immunosuppression.

The tumor microenvironment plays a fundamental role in B-cell lymphoma pathogenesis, largely by influencing survival and growth signalings and immune infiltration or blockade^64^. While disruption of follicular structures has long been recognized in B-cell lymphomas^65^, the spatial segregation of maturation states within tumors—such as FDC and TFH co-localizing with GC tumor states—may explain how different maturation states can differentiate within a tumor. The greater diversity of maturation states in FL compared to GCB DLBCL suggests that retention of follicular structures may better preserve the maturation process. With the success of CAR-T and bi-specific T-cell engagers (BiTEs) in relapsed or refractory DLBCL^66,67^, varying patterns of T-cell interactions across intratumor maturation states raise the question of whether immunotherapy should be tailored to the tumor’s maturation state composition, potentially requiring combination approaches to account for multiple spatial niches. Additionally, the diverse microenvironments around tumor maturation states encourage research into expanding the immunotherapy repertoire, including myeloid cells and non-malignant B cells.

The association of genetic variants with intratumor maturation states suggests that oncogenic mechanisms are linked to the differentiation process, with specific aberrations potentially more likely to lead to malignancy at different stages of B-cell maturation. Distinct expression patterns of genes with mutations between maturation states support this hypothesis. This could explain why oncogenic mechanisms vary across B-cell lymphoma subtypes. Extending this concept, ongoing differentiation trajectories may promote different survival and proliferation mechanisms within cells of the same tumor. Our recent work with joint single-cell DNA and RNA sequencing of FL and DLBCL validated this, showing that BCL2 variants were enriched in LZ compared to DZ states within the same tumors^56^. Characterizing the association between genetic aberrations and tumor maturation states across space and time would provide powerful insights into the relationship between cellular differentiation and clonal evolution.

Analogous to variation between organisms drives the evolution of species^68^, intratumor heterogeneity drives tumor evolution. We show that the differentiation trajectory of B cells in lymph nodes, B-cell maturation, is a major source of variation in B-cell lymphomas. Intratumor heterogeneity in maturation states blurs subtype boundaries, whereby individual DLBCL and FL tumors can contain multiple clinical subtypes. Gene regulatory networks and cellular interactions central to the B-cell maturation process are predominantly retained in malignancy, while intratumor maturation states occupy distinct spatial microenvironments and may undergo genetic variation. Tumor maturation state profiling revealed subgroups of FL, GCB DLBCL, and ABC DLBCL with striking differences in survival, driven by DZ and IgM memory states. Seeing cellular differentiation as a source of variation in cancer not only illuminates the origins of intratumor heterogeneity, but also sheds light on tumor evolution trajectories and sets the stage for treatment strategies against the emergence of resistant phenotypes.

## Acknowledgments

D.F. was supported by two projects from the German Federal Ministry of Education and Research (SIMONA under grant agreement number 031L0263A and SMART-CARE under grant agreement number 031L0212E), the European Research Council Synergy Grant DECODE under grant agreement No. 810296 and the PhD program of the European Molecular Biology Laboratory (EMBL). T.R. was supported by the German Federal Ministry of Education and Research (SIMONA under grant agreement number 031L0263A) and a physician-scientist fellowship of the Medical Faculty of Heidelberg University. M.A.B. was supported by a Career Development award from the International Myeloma Society (IMS). H.V. was supported by the German Federal Ministry of Education and Research (SIMONA under grant agreement number 031L0263A). N.L. was supported by the Heidelberg School of Oncology (HSO2) fellowship from the National Center for Tumor Diseases Heidelberg. O.W. is supported by an Else-Kröner Excellence Fellowship from the Else-Kröner-Fresenius Stiftung (Project-ID 2021_EKES.13) and the German Research Foundation (DFG, #WE4679/2-1). S.D. was supported by a grant from the Hairy Cell Leukemia Foundation, the Heidelberg Research Centre for Molecular Medicine (HRCMM), and an e:med junior group grant from the German Federal Ministry of Education and Research (BMBF). For the data management, we thank the Scientific Data Storage Heidelberg (SDS@hd) which is funded by the state of Baden-Wurttemberg and a DFG grant (INST 35/1314-1 FUGG). We thank Carolin Kolb (University Hospital Heidelberg), the EMBL Genomics Core Facility (Genecore), and the DKFZ Single-Cell Open Lab (scOpenLab) for their excellent technical assistance. Survival data was provided by the Molecular Mechanisms in Malignant Lymphoma (MMML) Network which received funding from the Deutsche Krebshilfe (DKH). We thank Sarah Kaspar and Felix Schneider at EMBL Data Science Internal Support for providing statistical advice. This manuscript was edited at Life Science Editors.

## Author Contributions

D.F., W.H., and S.D. conceptualized the project. D.F., T.R., and M.K. performed the single-cell experiments. D.F., T.R., J.M., L.C., P.M.B., and N.L. obtained clinical documentation. D.F., T.R., B.B., J.M., and L.L. performed single-cell data preprocessing and exploratory analyses. A.K. identified B-cell maturation states with flow cytometry and sorted them for RNA sequencing. DF. and B.B. analyzed the RNA sequencing data from sorted maturation states. D.F. characterized malignancy, transcriptional phenotypes, and B-cell maturation states in the CITE-Seq and 5’ BCR data. D.F. performed the maturation gene expression signature scoring. D.F. and A.M. analyzed transcription factor activity. M.A.B. performed the CODEX imaging, with preprocessing by M.A.B., H.V., and E.C. D.F., M.A.B., H.V., A.H., E.C., A.M., and F.C. characterized cell types in the CODEX data. D.F. developed and performed label transfer of B-cell maturation states from the CITE-Seq to CODEX data. A.H. performed cellular neighborhood analysis and characterized neighborhoods with D.F. E.C. and M.M.B. performed cellular interactions likelihood analysis. V.P. analyzed genetic variants in the DNA sequencing data. D.F. inferred and analyzed copy number variation in the single-cell data. R.S. supported the interpretation of the MMML data. W.H. and S.D. acquired most funding and resources. D.F. wrote the original draft which was initially reviewed by M.S., W.H., and S.D. All authors subsequently reviewed the paper.

## Disclosure of Conflicts of Interest

G.P.N. is a co-founder and stockholder of Akoya Biosciences, Inc. and inventor on patent US9909167 (On-slide staining by primer extension).

## Notes

### Summary of Updates

Adjustments to the text were made for clarification of the findings and their implications. Survival analysis (Figure 7) was added, highlighting distinct risk groups within diagnoses based on intratumor maturation state profiling.

