## Supplemental Data for "A single-cell multi-omic and spatial atlas of B-cell lymphomas reveals differentiation drives intratumor heterogeneity"

### Methods

#### Lymph node sample processing

The University of Heidelberg's Ethics Committee approved our study (S-254/2016), and we secured informed consent from every patient beforehand. We processed and froze patient lymph node (LN) samples for later analysis, following previously described methods[6]. To mitigate the influence of treatment-associated effects on tumor cells and their surroundings, we excluded samples from patients who had undergone allogeneic stem cell transplantation, CAR T-cell, or bispecific antibody therapy from the CITE-Seq cohort. Furthermore, we ensured all samples were collected at least three months post the termination of the most recent treatment to maintain the same control. We provide an overview of the sample composition in Supplemental Table 1.

#### Single-cell 3' RNA-seq and epitope expression profiling (CITE-Seq)

The cells were thawed, promptly washed to eliminate DMSO, and processed in groups of four to five that comprised a minimum of three distinct entities to avert entity-driven batch effects. A dead cell removal kit from Miltenyi Biotec was employed after thawing, aiming for cell viability of between 85% to 90%. Samples with less than 85% viability were not included. We then stained 5 x 10^5 live cells with a cocktail of oligonucleotide-linked antibodies (Supplemental Table 2) and left them to incubate at 4°C for 30 minutes. The cells were washed thrice with chilled washing buffer and centrifuged for five minutes at 4°C each time. Following this, cell count and viability were re-evaluated; samples falling below 85% viability were discarded. Subsequently, we prepared the bead-cell suspensions and carried out the synthesis of complementary DNA, single-cell gene expression, and the production of antibody-derived tag (ADT) libraries. For these steps, we used a Chromium single-cell v3.1 3' kit from 10x Genomics and followed the manufacturer's guidelines (Supplemental Table 1).

#### Single-cell 5' RNA-seq and B-cell receptor repertoire profiling

Apart from epitope staining, sample processing was identical to 3' scRNA-seq. The preparation of the bead-cell suspensions, synthesis of complementary DNA and single-cell gene expression, and BCR libraries were performed using a Chromium single-cell v2 5' and human BCR amplification kit (both 10x Genomics) according to the manufacturer's instructions. An overview of sample libraries is provided in Supplemental Table 1.

**Single-cell library sequencing and data processing**

We pooled the 3' gene expression and ADT libraries in a 3:1 ratio, targeting 40,000 reads (gene expression) and 15,000 reads per cell (ADT) respectively, and sequenced them on a NextSeq 500 (Illumina). 5' gene expression libraries were sequenced on a NextSeq 2000 (Illumina), aiming for 50,000 reads per cell. BCR libraries, sequenced on a NextSeq 500 (Illumina), were aimed at achieving a minimum of 5,000 reads per cell.

Post sequencing, we utilized the Cell Ranger software's (10x Genomics, v6.1.1) *cellranger mkfastq* function for demultiplexing and aligning raw base-call files to the reference genome (hg38). For 3' gene and epitope expression libraries, we used the *cellranger count* command on the resulting FASTQ files, while we used *cellranger multi* for 5' gene expression and BCR libraries. For the BCR libraries, we used the VDJ Ensembl reference (hg38, v5.0.0) as a reference. Unless specifically stated otherwise, we adhered to default settings for all functions.

**CITE-Seq data analysis**

The *Seurat* R package (v5.1.0) was used to perform data quality control, filtering, and normalization (log-based normalization for RNA and centered log-ratio transformation for ADT data). Gene counts per cell, ADT counts per cell, and percentages of mitochondrial reads were computed using the built-in functions. After quality control, we obtained data for 154,282 B cells with a median of 2,988 B cells per sample [140, 7,868] and a median of 6,887 transcript and 2,532 surface protein counts per cell. Principal component analysis[7], Louvain clustering[8], and UMAP[9] were performed for the transcriptome (RNA) and epitope (ADT) data independently. After mapping the CD3 and CD19 epitope expression, non-B-cell transcriptomic clusters and doublets were removed. We used the IntegrateData function of the Seurat package for data integration across the different preparation batches. For multimodal clustering, multi-omic factor analysis was performed with the *MOFA2* R package[3,10] (v1.8) based on the combined transcriptome and epitope data, and the resulting latent factors (n=30) were used as principal components.

**5' single-cell RNA-seq data and B-cell receptor profile analysis**

Transcriptomic analysis was performed using the R package Seurat (v4.1.0) as described in the *CITE-Seq data analysis* section above. B-cell receptor (BCR) clonotypes were added to the metadata from the *cellranger multi* output.

**Sorting of B-cell maturation states from reactive lymph nodes**

5 reactive lymph node and 2 tonsil samples were thawed as per a previously published protocol[6]. Using the BD FACSAria Fusion cell sorter equipped with the BD FACSDiva software v. 9.0 and FlowJo analysis software version 10.9.0, B cell maturation stated were identified and sorted with the gating strategy outlined in Supplemental Figure 1. Maturation states and their marker panels were sourced from previous studies which defined naïve[11,12], germinal center[13], memory[11,12], and plasma[14] states with flow cytometry:

Naïve B cells (Naïve): CD19+, CD20+, CD38 low, CD27-, IgD high
Germinal center dark zone B cells (DZ): CD19+, CD20+, CD38+, CD184+, CD83-
Germinal center light zone B cells (LZ): CD19+, CD20+, CD38+, CD184-, CD83+
IgM memory B cells (Mem IgM): CD19+, CD20+, CD38 low, CD27+, IgM+
IgG memory B cells (Mem IgG): CD19+, CD20+, CD38 low, CD27+, IgG+
Plasmablasts/plasma cells (Plasma): CD19+, CD20 low, CD38 high, CD27 high, IgD low

**RNA-seq of sorted maturation states**

RNA was isolated by the RNeasy Micro Kit (Qiagen, Hilden, Germany) and quantified with Bioanalyzer RNA 6000 pico assay (Agilent, Santa Clara, US). The libraries were generated with NuGENs Trio RNA-Seq System (NuGEN, Redwood City, California) for whole RNA and sequenced on an Illumina NextSeq2000 (Illumina, San Diego, US). Reads were trimmed with TrimGalore v0.6[15] and aligned with hisat2 v2.2.1[16]. The *DESeq2* R package[17] (v1.38.3) was used for differential gene expression analysis between maturation states. Default parameters were used unless otherwise specified. Gene symbols, as per the scRNA-seq datasets, were obtained from *Ensembl*[18] HGNC symbols.

**Characterization of B-cell maturation states in the reactive lymph node reference**

Clustering and differential expression analysis of the scRNA-seq data from B cells in the integrated reactive lymph node samples (8 samples, 16625 cells) were performed as described in the *Seurat Guided Clustering Tutorial* [19], with the clustering resolution parameter set to 1. Clusters were assigned to known B-cell maturation states based on their differential expression (in RNA and ADT features) of established markers of B-cell maturation states from several sources in the literature[2,20–26] (Supplemental Table 3).

**Classification of reactive lymph node maturation states in single-cell RNA-seq data**

The RNA-seq data from the sorted maturation states was filtered by the 2000 most variable RNA features in the reactive lymph node scRNA dataset (8 samples combined) and scaled. The resulting matrix was used as input for training a logistic regression model with 10×10-fold nested cross-validation for the classification of maturation states using the *nestedcv* package[27–30]. The resulting best-fit model was used to predict maturation states in the log-normalized and scaled scRNA-seq data from reactive lymph nodes. Predicted states were used to validate marker-based maturation state annotations in the CITE-Seq rLN reference dataset.

**Mapping of maturation states in all lymph node samples**

B-cell maturation states defined in the reactive lymph node reference were mapped to each tumor sample in the CITE-Seq and 5' scRNA-seq datasets using an anchor-based single-cell integration approach outlined in the *Seurat multimodal reference mapping* tutorial[31,32]. Log-normalized counts (without batch-effect correction to prevent bias introduced by sample integration) were used to find transfer anchors and project samples on the reference reductions - PCA (50 dimensions) and UMAP (2 dimensions).

**Isolation of malignant B cells**

Malignant B cells in tumor samples were identified based on immunoglobulin light chain restriction, whereby malignant (monoclonal) populations of cells are restricted to either the kappa or lambda immunoglobulin light chain and non-malignant B-cell populations (polyclonal) show mixed kappa and lambda light chain positivity[33]. As a minority of ADT counts may be present from ambient unbound antibodies during CITE-Seq library preparation, the proportion of total light chain counts of the kappa subtype (Kappa counts/(Kappa + Lambda counts)) per cell was used as a surrogate for binary positivity. Transcriptional B-cell clusters with an average kappa light chain proportion of >80% or <20% across all cells were considered malignant. Non-malignant B cells represented a median of 6% [0%, 94%] of all B cells in tumor samples.

**Maturation state gene expression signature scoring**

Maturation state gene expression signature scores were calculated by averaging the log-normalized counts for the 50 most differentially expressed genes (by fold-change) for each B-cell maturation state annotated in a published tonsil scRNA-seq dataset[2].

**Inference of transcription factor activity from single-cell RNA-sequencing data**

The pySCENIC[4] workflow was executed through a custom Snakemake pipeline. To infer the Gene Regulatory Network (GRN), we used the GRNBoost2 algorithm from the Arboreto[34] package with 10 perturbations. The analysis was performed on the raw scRNA-seq data. Transcription factor (TF) regulons were predicted using the human v9 motif collection from cisTarget (hg38__refseq-r80__10kb_up_and_down_tss.mc9nr.feather and hg38__refseq-r80__500bp_up_and_100bp_down_tss.mc9nr.feather databases). AUC scores per cell and GRNs were obtained for visualization and downstream analysis. For the final GRN reconstruction, only target genes occurring in more than 95% of the runs were considered. Differential expression (DE) analysis between B-cell maturation stages was conducted using Seurat's FindMarkers[35] function on the RNA assay, utilizing the MAST[5] method for DE analysis from single-cell data. DE genes between conditions in all cell populations were identified ($p.adj < 10e-16$ & $log2FC > 0.4$), and p-values were adjusted for multiple comparisons using the Benjamini-Hochberg[36] correction method. To determine differentially active TFs, we utilized the output of the SCENIC GRN, which provided TF activity at a single-cell level. Differentially active TFs were detected using Fisher's exact test to assess the enrichment of maturation stage-specific DE genes among all the TF target genes extracted from the SCENIC GRN ($p.adj < 0.05$).

**CODEX sample preparation**

Representative tumor or tumor-free lymph node areas were selected from archival FFPE tissue blocks belonging to 19 patients. This selection was made by the certified pathologists at the National Center for Tumor Diseases' Tissue Bank and the University Hospital Heidelberg's Institute of Pathology as previously described[37]. Two 4.5 mm cores per patient were incorporated into Tissue Microarrays (TMAs). TMA sections (4 µm) were affixed to Vectabond-precoated 25 x 25 mm coverslips, coated with paraffin, and stored for future staining.

#### Antibody conjugation, validation, and titration

We used the co-detection by indexing (CODEX) approach for multicolor immunofluorescence[38]. Antibodies utilized for CODEX experiments are summarized in Supplemental Table 2. We reduced purified, carrier-free antibodies with Tris(2-carboxyethyl)phosphine (TCEP) and conjugated them with maleimide-modified CODEX DNA oligonucleotides, procured from TriLink Biotechnologies. A board-certified pathologist supervised the evaluation of the conjugated antibodies in singleplex stains on tonsil and/or lymphoma tissue, comparing with online databases, immunohistochemical reference stains, and published literature. We validated staining patterns in multiplex experiments with positive and negative control antibodies and titrated the appropriate dilution of each antibody starting from 1:100 to optimize the signal-to-noise ratio.

#### Multiplex tissue staining and fixation

We deparaffinized, and rehydrated coverslips, and subjected them to heat-induced epitope retrieval at pH9 and 97°C for 10 minutes in a Lab Vision PT module. After blocking non-specific binding with CODEX FFPE blocking solution, we stained the coverslips overnight with the full antibody panel at the dilutions shown in Supplemental Table 2. Following staining, coverslips were fixed with 1.6% paraformaldehyde, methanol, and BS3 fixative, then stored in CODEX buffer S4 until imaging.

#### Multicycle imaging

We attached stained coverslips to custom acrylic plates and inserted them into a Keyence BZ-X710 inverted fluorescence microscope. We selected 7x7 fields of view and an appropriate number of z-planes (10-14) to capture the best focal plane across the imaging area. Multicycle imaging was performed using a CODEX microfluidics device. Post completion of multicycle imaging, coverslips were stained with hematoxylin/eosin, and the same areas were imaged in brightfield mode.

#### Image processing

We processed raw TIFF images using the RAPID pipeline[39] in Matlab with the default settings. Post-processing, images were concatenated to hyperstacks. Each tissue core was visually inspected for staining quality using ImageJ/Fiji.

**Cell segmentation and cell type annotation**

We segmented individual nuclei based on the Hoechst stain and quantified cellular marker expression levels using a modified version of the Mask R-CNN-based CellSeg software. A threshold based on the intensity of the nuclear markers Hoechst and DRAQ5 was used to exclude non-cellular events. Cells were then submitted to Leiden-based clustering using the scanpy Python package, and cluster annotations were assigned according to previously identified cell type marker profiles[37].

**CITE-Seq to CODEX B-cell maturation state label transfer**

B-cell maturation states in the CODEX data were classified sample-wise from annotations in the CITE-Seq data using shared features (n = 28) in the CITE-Seq and CODEX antibody panels (Supplemental Table 2). After selecting the shared features, CITE-Seq ADT counts and CODEX fluorescence intensities were subject to the same preprocessing steps of log-ratio normalization and scaling (z-scored) with *Seurat* v4. For each sample, a logistic regression classifier (*glmnet* package, 10×10-fold nested cross-validation)[29,30] was trained on the annotated CITE-seq data to classify B-cell maturation states. To prevent prediction bias toward majority classes, random sampling was performed to balance class distribution within the splits. For each sample, the resulting best-fit model (with the highest balanced accuracy on the outer folds) was used to predict B-cell maturation states in the sample's corresponding CODEX B-cell data. The median Pearson correlation coefficient between the samples' CITE-Seq and CODEX maturation state proportions was 0.91 ($p = 0.011$) (Supplemental Fig.8).

**Cellular neighborhood analysis**

We modified a previously described approach for neighborhood analysis[40]. For each cell, the 20 nearest neighbors were determined based on their Euclidean distance of the X and Y coordinates, thereby creating one 'window' of cells per individual cell. Next, we grouped these windows using k-means clustering according to the proportions of cell types within each window. We selected K=11 for the number of neighborhoods as we observed that higher values of k did not result in an improved biologically interpretable number of neighborhoods. Neighborhoods were annotated based on their biological function in normal lymph nodes or their enriched cell type(s)/state(s).

**Cellular interaction likelihood analysis**

Spatial graph representations of immediately neighboring cells were constructed based on Delaunay triangulation between centroid coordinates using the *scipy.spatial* Python package[41]. To compute pairwise association strengths between clusters, relative frequencies were computed using the following metric:

$$\frac{Nij \ \times Nt}{Ni \times Nj}$$

in which Nij is equal to the total number of edges between clusters i and j, Nt the total number of edges in the sample, and Ni and Nj the total degrees of clusters i and j respectively[40]. Computed association strengths were calculated separately for each disease entity, between B-cell states and other cell types.

**DNA sequencing**

DNA was fragmented (Covaris sonication) to 250 bp and further purified using Agentcourt AMPure XP beads (Beckman Coulter). Size-selected DNA was then ligated to adaptors during library preparation. Each library was quantified using qPCR and analyzed for quality after fragmentation and library preparation based on library yield and size on an Agilent Bioanalyzer. The sample $MZL_2$ failed at the library preparation stage. Finally, libraries were enriched for genes using the Sure Select XT Target Enrichment System for Illumina Paired-End Multiplexed Sequencing and each capture pool was sequenced at 300-400x. A list of captured regions is included in Supplemental Table 4.

Pooled samples were demultiplexed using a custom demultiplexing tool. Read pairs were aligned to the hg19 reference sequence using the Burrows-Wheeler Aligner[42], and data were sorted and duplicate-marked using Picard tools (version 2.23.3)[43]. All steps were performed within the bcbio-nextgen toolkit (version 1.2.9)[44].

The minimum quality criterion was 80% of target bases having > 30x sequencing coverage. Cases with 60-79% of target bases with > 30x sequencing coverage were also included if target bases not covered were < 1%. Cases with target bases covered 30x < 60% or cases with target bases covered 30x between 60-80% and target bases not covered > 1% were excluded.

This was achieved for all sequenced samples (Supplemental Fig. 11). Metrics were collected using Picard tools (version 2.23.3)[43]. For detailed QC metrics see Supplemental Table 5.

**Variant Analysis**

Mutation analysis for single nucleotide variants (SNV) and Insertions and Deletions (InDels) was performed using MuTect2[45] (GATK v4.1.9.0)[46] and annotated by Funcotator[47] (GATK v4.1.9.0). A panel-of-normals (PON) filter was generated using samples annotated as rLN and a panel of normal from the 1000 Genomes Project[48]. Variants were included in the PON if present in two or more normal samples.

Non-silent variants (Missense_Mutation, Nonsense_Mutation, Nonstop_Mutation, Splice_Site, Translation_Start_Site) resulting from BestEffect Funcotator annotation (dataSources.v1.6) at a variant allele frequency of > 10% are kept for further investigations. Germline polymorphisms and sequencing artifacts were excluded by comparison with the panel-of-normals and with the gnomAD database[49]. Known germline polymorphisms from the Exome Sequencing Project[50] and dbSNP[51] databases were excluded. An overview of the somatic variants identified is depicted in the Supplemental Fig. 11.

**Inference of copy number variation from single-cell RNA-sequencing data**

Copy number variants (CNVs) and ploidy were inferred from single-cell RNA-sequencing count data in each sample using the copykat R package as per the package vignette[53]. A cell filtering threshold of 5 genes per chromosome and a minimal segmentation window size of 25 genes was used. Copy number variation (Euclidean distance) was determined at a resolution of 5MB chromosomal segments, which was added as a new assay to CITE-Seq Seurat objects for each sample for visualization of copy number variants across intratumor maturation states.

**Analysis of IGHV variants in tumor maturation states**

Single-cell DNA and RNA sequencing (SDR-seq) data was obtained from Lindenhofer et al., 2024[54], with the genetic variants and maturation states identified as described in the publication. Genetic variants detected in immunoglobulin heavy chain variable region (IGHV) genes that were exclusive to the B cells were identified and their genotype was plotted for 50 randomly selected cells per state.

#### Deconvolution of tumor cell type composition from bulk microarray data

The CITE-Seq dataset (51 samples) was downsampled to 100 cells per annotated cell type, including the B-cell maturation states we describe here and the immune cell populations described in Roider at al. 2024 [55]. Using the CIBERSORTx[56] web interface, we generated a signature matrix from the downsampled single-cell gene expression counts. Bulk gene expression data (Affymetrix HG-U133A GeneChip microarrays) from 430 DLBCL (142 GCB, 133 ABC, 97 unclassified, 58 double-hit), 145 FL (114 BCL2-break positive, 29 BCL2-break negative, 2 BCL-break unknown), and 48 mixed FL/DLBCL (FL grade 3B) tumors was obtained from Loeffler-Wirth et al. 2019[57]. Microarray probes were converted to gene symbols with the annotate (v1.82.0) and hgu133plus2.db (v3.13.0) R packages and gene expression data were counts-per-million-normalized to create a mixture file. Using the signature matrix file described above, we imputed fractions of cell types from our CITE-Seq dataset in the bulk gene expression data with CIBERSORTx[56]. Batch correction (S-mode) was enabled and quantile normalization was disabled, with 100 permutations for significance analysis. Default CIBERSORTx parameters were used unless otherwise specified.

#### Survival analysis with tumor cell type composition

After filtering for samples with follow-up survival data from the Loeffler-Wirth et al. 2019[57] dataset described above, 507 tumors remained (119 DLBCL GCB, 102 DLBCL ABC, 79 unclassified DLBCL, 44 double-hit DLBCL, 94 BCL2-break positive FL, 25 BCL2-break negative FL, 2 BCL-break unknown FL, and 42 mixed FL/DLBCL (FL grade 3B)). Cell type proportions for each sample were divided by the mean proportion of the respective cell type across samples. Cox proportional hazard ratios for overall survival were computed independently for each maturation state using the survival R package (v3.7.0)[58]. P-values were adjusted for multiple hypothesis testing using the Benjamini-Hochberg method[36]. A multivariate Cox proportional hazard model was generated to predict survival based on the proportions of all cell types in each tumor using the glmnet R package (v4.1.8)[59,60]. Nested cross-validation was performed, whereby the optimal lambda was determined from the inner folds (n=5), and the concordance index (C-index) was calculated on the outer folds (n = 5). The final model was used to segregate samples into high, medium, and low-risk groups of equal size based on their predicted survival. Kaplan-Meier survival curves were plotted with the survminer R package (v0.4.9)[61], on which p-values are reported for log-rank test for differences in survival between groups. Optimal split points on cell type proportions for creating distinct survival groups were determined with the maxstat R package (v0.7-25) using the Log Rank test.

21. Seifert, M., Scholtysik, R. & Küppers, R. Origin and Pathogenesis of B Cell

Lymphomas. in *Lymphoma: Methods and Protocols* (ed. Küppers, R.) 1–33 (Springer New York, New York, NY, 2019). doi:10.1007/978-1-4939-9151-8_1.

32. Multimodal reference mapping. https://satijalab.org/seurat/articles/multimodal_reference_mapping.html.
33. Ratech, H. & Litwin, S. Surface immunoglobulin light chain restriction in B-cell non-Hodgkin's malignant lymphomas. *Am. J. Clin. Pathol.* **91**, 583–586 (1989).
34. Moerman, T. *et al.* GRNBoost2 and Arboreto: efficient and scalable inference of gene regulatory networks. *Bioinformatics* **35**, 2159–2161 (2019).
35. Hao, Y. *et al.* Integrated analysis of multimodal single-cell data. *Cell* **184**, 3573–3587.e29 (2021).
36. Benjamini, Y. & Hochberg, Y. Controlling the False Discovery Rate: A Practical and Powerful Approach to Multiple Testing. *J. R. Stat. Soc. Series B Stat. Methodol.* **57**, 289–300 (1995).
37. Roider, T. *et al.* Multimodal and spatially resolved profiling identifies distinct patterns of T-cell infiltration in nodal B-cell lymphoma entities. *bioRxiv* 2022.11.04.514366 (2022) doi:10.1101/2022.11.04.514366.
38. Black, S. *et al.* CODEX multiplexed tissue imaging with DNA-conjugated antibodies. *Nat. Protoc.* **16**, 3802–3835 (2021).
39. Lu, G. *et al.* A real-time GPU-accelerated parallelized image processor for large-scale multiplexed fluorescence microscopy data. *Front. Immunol.* **13**, 981825 (2022).
40. Schürch, C. M. *et al.* Coordinated Cellular Neighborhoods Orchestrate Antitumoral Immunity at the Colorectal Cancer Invasive Front. *Cell* **182**, 1341–1359.e19 (2020).
41. Virtanen, P. *et al.* SciPy 1.0: fundamental algorithms for scientific computing in Python. *Nat. Methods* **17**, 261–272 (2020).
42. Li, H. & Durbin, R. Fast and accurate short read alignment with Burrows–Wheeler transform. *Bioinformatics* **25**, 1754–1760 (2009).
43. Wysoker, A., Tibbetts, K. & Fennell, T. Picard tools version 1.90. *Jhpsn.*

44. Guimera, R. V. bcbio-nextgen: Automated, distributed next-gen sequencing pipeline. *EMBnet J.* **17**, 30 (2012).

45. Benjamin, D. *et al.* Calling Somatic SNVs and Indels with Mutect2. *bioRxiv* 861054 (2019) doi:10.1101/861054.

46. *Gatk: Official Code Repository for GATK Versions 4 and up*. (Github).

47. Funcotator Information and Tutorial. *GATK* https://gatk.broadinstitute.org/hc/en-us/articles/360035889931-Funcotator-Information-and-Tutorial.

55. Roider, T. *et al.* Multimodal and spatially resolved profiling identifies distinct patterns of

T cell infiltration in nodal B cell lymphoma entities. *Nat. Cell Biol.* (2024) doi:10.1038/s41556-024-01358-2.

56. Newman, A. M. *et al.* Determining cell type abundance and expression from bulk tissues with digital cytometry. *Nat. Biotechnol.* **37**, 773–782 (2019).
57. Loeffler-Wirth, H. *et al.* A modular transcriptome map of mature B cell lymphomas. *Genome Med.* **11**, 27 (2019).
58. Therneau, T. M. & Grambsch, P. M. *Modeling Survival Data: Extending the Cox Model*. (Springer Science & Business Media, 2013).
59. Friedman, J., Hastie, T. & Tibshirani, R. Regularization Paths for Generalized Linear Models via Coordinate Descent. *J. Stat. Softw.* **33**, 1–22 (2010).
60. Simon, N., Friedman, J., Hastie, T. & Tibshirani, R. Regularization Paths for Cox's Proportional Hazards Model via Coordinate Descent. *J. Stat. Softw.* **39**, 1–13 (2011).
61. Kassambara, A., Kosinski, M., Biecek, P. & Fabian, S. survminer: Drawing Survival Curves using 'ggplot2'. 2019. *R package version 0. 4* **4**, (2021).

### Tables

#### Supplemental Table 1: CITE-seq sample overview

| *Sample ID* | *Run* | *Entity* | *Age* | *Sex* | *Status* | *Ann-Arbor Stage* |
|---|---|---|---|---|---|---|
| *ABC1* | 15 | DLBCL, non-GCB | 67 | M | Initial diagnosis | IIIB |
| *ABC2* | 6 | DLBCL, non-GCB | 63 | M | Initial diagnosis | III |
| *ABC3* | 8 | DLBCL, non-GCB | 80 | M | Initial diagnosis | IIIB |
| *ABC4* | 8 | DLBCL, non-GCB | 61 | F | Initial diagnosis | IIIB |
| *ABC5* | 3 | DLBCL, non-GCB | 38 | F | Relapse | IVA |
| *ABC6* | 8 | DLBCL, non-GCB | 69 | M | Relapse | IAE |
| *ABC7* | 6 | DLBCL, non-GCB | 54 | M | Relapse | IIIA |
| *FL1* | 10 | FL | 64 | F | Initial diagnosis | IVB |
| *FL10* | 2 | FL | 71 | M | Relapse | IVA |
| *FL11* | 11 | FL | 74 | M | Relapse | IVA |
| *FL12* | 9 | FL | 66 | M | Relapse | IVA |
| *FL2* | 16 | FL | 78 | M | Initial diagnosis | IIIA |
| *FL3* | 3 | FL | 67 | F | Initial diagnosis | IVA |
| *FL4* | 1 | FL | 33 | M | Initial diagnosis | IVB |
| *FL5* | 2 | FL | 54 | M | Relapse | IIIA |
| *FL6* | 1 | FL | 67 | M | Relapse | IVA |
| *FL7* | 10 | FL | 59 | F | Relapse | IIIA |
| *FL8* | 8 | FL | 42 | M | Relapse | IIA |
| *FL9* | 12 | FL | 72 | F | Relapse | IIIA |
| *GCB1* | 16 | DLBCL, GCB | 52 | F | Relapse | IIIAE |
| *GCB2* | 5 | DLBCL, GCB | 84 | F | Initial diagnosis | IVA |
| *GCB3* | 4 | DLBCL, GCB | 57 | M | Initial diagnosis | IIIA |
| *GCB4* | 13 | DLBCL, GCB | 45 | M | Relapse | IVB |
| *GCB5* | 15 | DLBCL, GCB | 77 | M | Relapse | IIB |
| *MCL1* | 4 | MCL | 63 | M | Initial diagnosis | IVB |
| *MCL2* | 7 | MCL | 77 | M | Initial diagnosis | IVA |
| *MCL3* | 3 | MCL | 68 | M | Initial diagnosis | IVB |

| | | | | | | |
|---|---|---|---|---|---|---|
| *MCL4* | 9 | MCL | 50 | M | Initial diagnosis | IVB |
| *MCL5* | 10 | MCL | 69 | M | Initial diagnosis | IVB |
| *MCL6* | 15 | MCL | 61 | M | Relapse | IVA |
| *MCL7* | 12 | MCL | 62 | M | Relapse | IVA |
| *MCL8* | 11 | MCL | 72 | M | Relapse | IVA |
| *MZL1* | 4 | MZL | 49 | M | Initial diagnosis | IVA |
| *MZL10* | 11 | MZL | 78 | M | Relapse | IAE |
| *MZL11* | 9 | MZL | 63 | M | Relapse | IIIA |
| *MZL2* | 7 | MZL | 58 | F | Initial diagnosis | IV |
| *MZL3* | 3 | MZL | 49 | F | Initial diagnosis | IVB |
| *MZL4* | 14 | MZL | 34 | F | Initial diagnosis | IVA |
| *MZL5* | 16 | MZL | 71 | F | Initial diagnosis | IA |
| *MZL6* | 14 | MZL | 59 | F | Relapse | IIA |
| *MZL7* | 6 | MZL | 82 | M | Relapse | IV |
| *MZL8* | 5 | MZL | 52 | F | Relapse | IVA |
| *MZL9* | 16 | MZL | 54 | F | Relapse | IVA |
| *rLN1* | 7 | rLN | 73 | F | n / A | n / A |
| *rLN2* | 10 | rLN | 35 | F | n / A | n / A |
| *rLN3* | 6 | rLN | 33 | F | n / A | n / A |
| *rLN4* | 11 | rLN | 51 | M | n / A | n / A |
| *rLN5* | 14 | rLN | 20 | M | n / A | n / A |
| *rLN6* | 4 | rLN | 46 | F | n / A | n / A |
| *rLN7* | 7 | rLN | 23 | F | n / A | n / A |
| *rLN8* | 9 | rLN | 61 | M | n / A | n / A |

M = Male
F = Female
rLN = reactive lymph node
MCL = mantle cell lymphoma
FL = follicular lymphoma
DLBCL, (non-)GCB = diffuse large B-cell lymphoma, (non-)germinal center
MZL = marginal zone lymphoma

#### Supplemental Table 2: CITE-Seq antibody panel

| Specificity | Alternative | Clone | Isotype | Catalogue | BarcodeNo | Catalog PE | Titration | Concentration | Amount Used |
|---|---|---|---|---|---|---|---|---|---|
| CD10 | | HI10a | Mouse IgG1, κ | 312231 | 0062 | 312203 | 1.25 µl | 100 µg/ml | 125 ng |
| CD103 | | Ber-ACT8 | Mouse IgG1, κ | 350231 | 0145 | 350205 | 1.25 µl | 12 µg/ml | 15 ng |
| CD11b | | ICRF44 | Mouse IgG1, κ | 301353 | 0161 | | | | |
| CD11c | | S-HCL-3 | Mouse IgG2b, κ | 371519 | 0053 | 371504 | 0.63 µl | 50 µg/ml | 31 ng |
| CD127 | IL7R | A019D5 | Mouse IgG1, κ | 351352 | 0390 | 351303 | 0.63 µl | 100 µg/ml | 63 ng |
| CD134 | | Ber-ACT35 | Mouse IgG1, κ | 350033 | 0158 | 350003 | 0.63 µl | 200 µg/ml | 125 ng |
| CD137 | | 4B4-1 | Mouse IgG1, κ | 309835 | 0355 | 309803 | 1.25 µl | 100 µg/ml | 125 ng |
| CD150 | SLAM | A12 (7D4) | Mouse IgG1, κ | 306313 | 0870 | | | | |
| CD152 | CTLA4 | BNI3 | Mouse IgG2a, κ | 369619 | 0151 | 369603 | 0.63 µl | 200 µg/ml | 125 ng |
| CD16 | | 3G8 | Mouse IgG1, κ | 302061 | 0083 | 302007 | 0.63 µl | 100 µg/ml | 63 ng |
| CD161 | | HP-3G10 | Mouse IgG1, κ | 339945 | 0149 | 339903 | 1.25 µl | 120 µg/ml | 150 ng |
| CD183 | CXCR3 | G025H7 | Mouse IgG1, κ | 353745 | 0140 | 353705 | 0.63 µl | 100 µg/ml | 63 ng |
| CD184 | CXCR4 | 12G5 | Mouse IgG2a, κ | 306531 | 0366 | | | | |
| CD185 | CXCR5 | J252D4 | Mouse IgG1, κ | 356937 | 0144 | 356903 | 1.25 µl | 100 µg/ml | 125 ng |
| CD19 | | HIB19 | Mouse IgG1, κ | 302259 | 0050 | 302207 | 0.63 µl | 50 µg/ml | 31 ng |
| CD194 | CCR4 | L291H4 | Mouse IgG1, κ | 359423 | 0071 | 359411 | 0.63 µl | 50 µg/ml | 31 ng |
| CD195 | CCR5 | J418F1 | Rat IgG2b, κ | 359135 | 0141 | 359105 | 1.25 µl | 100 µg/ml | 125 ng |
| CD197 | CCR7 | G043H7 | Mouse IgG2a, κ | 353247 | 0148 | 353203 | 2.50 µl | 160 µg/ml | 400 ng |

| | | | | | | | | | |
|---|---|---|---|---|---|---|---|---|---|
| CD2 | | TS1/8 | Mouse IgG1, κ | 309229 | 0367 | | | | |
| CD20 | | 2H7 | Mouse IgG2b, κ | 302359 | 0100 | 302305 | 2.50 µl | 25 µg/ml | 63 ng |
| CD200 | - | OX-104 | | na | na | 329206 | 0.63 µl | 200 µg/ml | 125 ng |
| CD21 | | Bu32 | Mouse IgG1, κ | 354915 | 0181 | 354903 | 0.63 µl | 50 µg/ml | 31 ng |
| CD22 | | S-HCL-1 | Mouse IgG2b, κ | 363514 | 0393 | 363503 | 0.63 µl | 100 µg/ml | 63 ng |
| CD223 | LAG3 | 11C3C65 | Mouse IgG1, κ | 369333 | 0152 | 369305 | 1.25 µl | 100 µg/ml | 125 ng |
| CD23 | | EBVCS-5 | Mouse IgG1, κ | 338523 | 0897 | 338507 | 2.50 µl | 50 µg/ml | 125 ng |
| CD24 | | ML5 | Mouse IgG2a, κ | 311137 | 0180 | 311105 | 1.25 µl | 200 µg/ml | 250 ng |
| CD244 | | C1.7 | Mouse IgG1, κ | 329527 | 0189 | 329507 | 0.63 µl | 50 µg/ml | 31 ng |
| CD25 | IL2RA | BC96 | Mouse IgG1, κ | 302643 | 0085 | 302605 | 1.25 µl | 50 µg/ml | 63 ng |
| CD27 | | O323 | Mouse IgG1, κ | 302847 | 0154 | 302807 | 0.63 µl | 100 µg/ml | 63 ng |
| CD273 | PDL2 | 24F.10C12 | Mouse IgG2a, κ | 329619 | 0008 | 329605 | 0.63 µl | 100 µg/ml | 63 ng |
| CD274 | PDL1 | 29E.2A3 | Mouse IgG2b, κ | 329743 | 0007 | 329705 | 0.63 µl | 400 µg/ml | 250 ng |
| CD278 | ICOS | C398.4A | Armenian Hamster IgG | 313555 | 0171 | 313507 | 0.63 µl | 200 µg/ml | 125 ng |
| CD279 | PD1 | EH12.2H7 | Mouse IgG1, κ | 329955 | 0088 | 329905 | 1.25 µl | 50 µg/ml | 63 ng |
| CD28 | | CD28.2 | Mouse IgG1, κ | 302955 | 0386 | 302907 | 1.25 µl | 100 µg/ml | 125 ng |
| CD29 | part of VLA-4 | TS2/16 | Mouse IgG1, κ | 303027 | 0369 | | | 50 µg/ml | |
| CD3 | | UCHT1 | Mouse IgG1, κ | 300475 | 0034 | 300407 | 0.63 µl | 100 µg/ml | 63 ng |
| CD31 | | WM59 | | 303137 | 0124 | 303105 | 0.63 µl | 100 µg/ml | 63 ng |
| CD32 | | FUN-2 | Mouse IgG2b, κ | 303223 | 0142 | 303205 | 0.63 µl | 50 µg/ml | 31 ng |

| | | | | | | | | | |
|---|---|---|---|---|---|---|---|---|---|
| CD357 | GITR | 108-17 | Mouse IgG2a, κ | 371225 | 0360 | 371203 | 2.50 µl | 100 µg/ml | 250 ng |
| CD366 | TIM-3 | F38-2E2 | Mouse IgG1, κ | 345047 | 0169 | 345005 | 1.25 µl | 100 µg/ml | 125 ng |
| CD38 | | HIT2 | Mouse IgG1, κ | 303541 | 0389 | 303505 | 1.25 µl | 100 µg/ml | 125 ng |
| CD39 | | A1 | Mouse IgG1, κ | 328233 | 0176 | 328207 | 0.63 µl | 50 µg/ml | 31 ng |
| CD4 | | RPA-T4 | Mouse IgG1, κ | 300563 | 0072 | 300507 | 0.63 µl | 100 µg/ml | 63 ng |
| CD43 | | CD43-10G7 | Mouse IgG1, κ | 343209 | 0357 | 343203 | 0.63 µl | 400 µg/ml | 250 ng |
| CD44 | | IM7 | Rat IgG2b, κ | 103045 | 0073 | 103023 | 0.63 µl | 50 µg/ml | 31 ng |
| CD45 | | HI30 | Mouse IgG1, κ | 304064 | 0391 | 304007 | 1.25 µl | 10 µg/ml | 13 ng |
| CD45RA | | HI100 | Mouse IgG2b, κ | 304157 | 0063 | 304107 | 0.63 µl | 12 µg/ml | 8 ng |
| CD45RO | | UCHL1 | Mouse IgG2a, κ | 304255 | 0087 | 304205 | 0.63 µl | 40 µg/ml | 25 ng |
| CD47 | | CC2C6 | Mouse IgG1, κ | 323129 | 0026 | 323108 | 0.63 µl | 80 µg/ml | 50 ng |
| CD48 | | BJ40 | Mouse IgG1, κ | 336709 | 0029 | 336707 | 1.25 µl | 200 µg/ml | 250 ng |
| CD5 | | UCHT2 | Mouse IgG1, κ | 300635 | 0138 | 300607 | 0.63 µl | 100 µg/ml | 63 ng |
| CD56 | | QA17A16 | Mouse IgG1, κ | 392421 | 0084 | 392403 | 0.63 µl | 80 µg/ml | 50 ng |
| CD57 | | QA17A04 | Mouse IgG1, κ | 393319 | 0168 | 393307 | 0.63 µl | 200 µg/ml | 126 ng |
| CD62L | | DREG-56 | Mouse IgG1, κ | 304847 | 0147 | 304805 | 1.25 µl | 25 µg/ml | 31 ng |
| CD69 | | FN50 | Mouse IgG1, κ | 310947 | 0146 | 310905 | 1.25 µl | 50 µg/ml | 63 ng |
| CD7 | | CD7-6B7 | Mouse IgG2a, κ | 343123 | 0066 | 343105 | 0.63 µl | 100 µg/ml | 63 ng |
| CD70 | | 113-16 | Mouse IgG1 | 355117 | 0027 | 355103 | 1.25 µl | 200 µg/ml | 250 ng |
| CD73 | | AD2 | Mouse IgG1, κ | 344029 | 0577 | 344003 | | 400 µg/ml | |

| | | | | | | | | | |
|---|---|---|---|---|---|---|---|---|---|
| CD79b | | CB3-1 | Mouse IgG1, κ | 341415 | 0187 | 341404 | 1.25 µl | 50 µg/ml | 63 ng |
| CD86 | B7-2 | IT2.2 | Mouse IgG2b, κ | 305443 | 0006 | | | | |
| CD8a | | RPA-T8 | Mouse IgG1, κ | 301067 | 0080 | 301007 | 0.63 µl | 100 µg/ml | 63 ng |
| CD95 | Fas | DX2 | Mouse IgG1, κ | 305649 | 0156 | 305607 | 0.63 µl | 100 µg/ml | 63 ng |
| Isotype Ctrl | | MOPC-21 | Mouse IgG1, κ | 400199 | 0090 | | | | |
| Isotype Ctrl | | MPC-11 | Mouse IgG2b, κ | 400373 | 0092 | | | | |
| Isotype Ctrl | | MOPC-173 | Mouse IgG2a, κ | 400285 | 0091 | | | | |
| Isotype Ctrl | | RTK4530 | Rat IgG2b, κ | 400673 | 0095 | | | | |
| Isotype Ctrl | | HTK888 | Armenian Hamster IgG | 400973 | 0241 | | | | |
| Kappa | | MHK-49 | Mouse IgG1, κ | 316531 | 0894 | 316507 | 1.25 µl | 20 µg/ml | 25 ng |
| KLRG1/MAFA | | SA231A2 | Mouse IgG2a, κ | 367721 | 0153 | 367711 | 0.63 µl | 100 µg/ml | 63 ng |
| Lambda | | MHL-38 | Mouse IgG2a, κ | 316627 | 0898 | 316607 | 1.25 µl | 20 µg/ml | 25 ng |
| TIGIT | | A15153G | Mouse IgG2a, κ | 372725 | 0089 | 372703 | 1.25 µl | 25 µg/ml | 31 ng |

**Supplemental Table 3: FACS antibody panel**

| *Specificity* | *Species* | *Fluorochrome* | *Clone* | *Supplier* | *Catalogue* |
|---|---|---|---|---|---|
| *anti-IgM* | human | BUV-395 | G20-127 | BD Biosciences | 563903 |
| *anti-CD10* | human | FITC | HI10a | BD Biosciences | 332775 |
| *anti-IgD* | human | PerCP-Cy5.5 | IA6-2 | BD Biosciences | 561315 |
| *anti-CD23* | human | APC | M-L233 | BD Biosciences | 558690 |
| *anti-CD38* | human | APC-R700 | HIT2 | BD Biosciences | 561979 |
| *anti-CD24* | human | BV421 | ML5 | BD Biosciences | 562789 |
| *anti-CD5* | human | BV510 | UCHT2 | BD Biosciences | 563381 |
| *anti-CD20* | human | BV650 | 2H7 | BD Biosciences | 563780 |
| *anti-CD21* | human | PE | B-ly4 | BD Biosciences | 555422 |
| *anti-IgM* | human | PE-CF594 | G20-127 | BD Biosciences | 562539 |
| *anti-CD27* | human | PE-Cy7 | M-T271 | BD Biosciences | 560609 |
| *anti-CD184* | human | BV421 | 12G5 | BD Biosciences | 566282 |
| *anti-CD83* | human | PE | HB15e | BD Biosciences | 550634 |
| *anti-Ig κ Light Chain* | human | PE-Cy7 | G20-193 | BD Biosciences | 561328 |
| *anti-Ig λ Light Chain* | human | PE | JDC-12 | BD Biosciences | 555797 |
| *anti-CD3* | human | PE | HIT3a | BD Biosciences | 555340 |
| *anti-CD19* | human | APC-Cy7 | SJ25C1 | BD Biosciences | 557791 |
| *anti-CD45R/B220* | mouse | PE | RA3-6B2 | BD Biosciences | 553090 |
| *anti-CD45RB* | human | FITC | MEM-55 | Biolegend | 310206 |

#### Supplemental Table 4: CODEX antibody panel

| *Target* | *Alternative* | *Clone* | *Supplier* | *Catalogue* | *CODEX oligo* | *Dilution* | *Exposure [ms]* | *Cycle* |
|---|---|---|---|---|---|---|---|---|
| *Blank* | | | | | | | | 1 |
| *BCL6* | | K112-91 | BD Biosciences | 561520 | 79 | 1:25 | 500 | 2 |
| *GATA3* | | L50-823 | Cell Marque | custom | 2 | 1:50 | 500 | 3 |
| *CD185* | CXCR5 | D6L3C | Cell Signaling Technology | custom | 69 | 1:100 | 500 | 4 |
| *Tbet* | | D6N8B | Cell Signaling Technology | custom | 68 | 1:100 | 500 | 5 |
| *CD62L* | | B-8 | Santa Cruz Biotechnology | custom | 38 | 1:400 | 250 | 6 |
| *FoxP3* | | 236A/E7 | Invitrogen | 14-4777-82 | 61 | 1:100 | 500 | 7 |
| *CD163* | | EDHu-1 | Novus Biologicals | NB110-40686 | 59 | 1:50 | 500 | 8 |
| *Ki67* | | B56 | BD Biosciences | 556003 | 6 | 1:200 | 333 | 9 |
| *CD366* | TIM3 | polyclonal | Novus Biologicals | AF2365 | 44 | 1:100 | 500 | 10 |
| *PAX5* | | D7H5X | Cell Signaling Technology | custom | 66 | 1:200 | 200 | 11 |
| *CD134* | | Ber-ACT35 | Biolegend | 350002 | 75 | 1:100 | 500 | 12 |
| *IL10* | | polyclonal | R&D Systems | AF-217-NA | 67 | 1:100 | 500 | 13 |
| *CD5* | | vC5/473 + CD5/54/F6 | Novus Biologicals | NBP2-34583 | 25 | 1:50 | 500 | 14 |
| *CD206* | | MM0820-48L31 | Abcam | n/a | 55 | 1:200 | 500 | 15 |
| *CD25* | IL2RA | 4C9 | Cell Marque | custom | 24 | 1:200 | 500 | 16 |
| *CD16* | | D1N9L | Cell Signaling Technology | custom | 60 | 1:50 | 500 | 17 |
| *CD152* | CTLA4 | BSB-88 | BioSB | BSB 2885 (ASR) | 30 | 1:25 | 500 | 18 |
| *CD79a* | | HM47 | Biolegend | 333502 | 46 | 1:200 | 250 | 19 |

| | | | | | | | | |
|---|---|---|---|---|---|---|---|---|
| *CD57* | | HNK-1 | Biolegend | 359602 | 29 | 1:50 | 500 | 20 |
| *CD34* | | QBEnd/10 | Novus Biologicals | NBP2-34713 | 11 | 1:50 | 500 | 21 |
| *CXCL13* | | polyclonal | Novus | AF801 | 41 | 1:200 | 500 | 22 |
| *CD21* | | SP186 | Abcam | ab240987 | 15 | 1:100 | 500 | 23 |
| *CD7* | | MRQ56 | Cell Marque | custom | 63 | 1:100 | 500 | 24 |
| *Podoplanin* | | D2-40 | Biolegend | 916606 | 32 | 1:200 | 500 | 25 |
| *CD279* | PD1 | D4W2J | Cell Signaling Technology | custom | 23 | 1:50 | 500 | 26 |
| *HLA-DR* | | EPR3692 | Abcam | ab209968 | 65 | 1:100 | 333 | 27 |
| *CD223* | LAG3 | D2G4O | Cell Signaling Technology | custom | 42 | 1:25 | 500 | 28 |
| *CD20* | | rIGEL/773 | Novus Biologicals | NBP2-54591 | 48 | 1:200 | 250 | 29 |
| *CD56* | | MRQ-42 | Cell Marque | custom | 58 | 1:100 | 250 | 30 |
| *CD45RO* | | UCH-L1 | Santa Cruz Biotechnology | custom | 5 | 1:50 | 500 | 31 |
| *CD278* | ICOS | D1K2T | Cell Signaling Technology | custom | 74 | 1:200 | 500 | 32 |
| *CD90* | | EPR3132 | Abcam | ab181885 | 57 | 1:150 | 500 | 33 |
| *CD4* | | EPR6855 | Abcam | ab181724 | 20 | 1:100 | 500 | 34 |
| *CD11c* | | EP1347Y | Abcam | ab216655 | 49 | 1:200 | 333 | 35 |
| *CD3* | | MRQ-39 | Cell Marque | custom | 33 | 1:50 | 500 | 36 |
| *CD68* | | KP-1 | BioLegend | 916104 | 62 | 1:200 | 250 | 37 |
| *CD69* | | EPR21814 | Abcam | ab234512 | 36 | 1:500 | 250 | 38 |
| *CD14* | | EPR3653 | Abcam | ab226121 | 7 | 1:300 | 250 | 39 |
| *CD8* | | C8/144B | Cell Marque | custom | 8 | 1:100 | 250 | 40 |

| | | | | | | | |
|---|---|---|---|---|---|---|---|
| *Kappa light chain* | L1C1 | Cell Marque | custom | 70 | 1:100 | 200 | 41 |
| *CD45RA* | HI100 | Biolegend | 304102 | 21 | 1:200 | 125 | 42 |
| *CD11b* | EPR1344 | Abcam | ab209970 | 28 | 1:200 | 167 | 43 |
| *Granzyme B* | EPR20129-217 | Abcam | ab219803 | 81 | 1:200 | 250 | 44 |
| *CD31* | C31.3 + C31.7 + C31.10 | Novus Biologicals | NBP2-47785 | 51 | 1:200 | 167 | 45 |
| *CD45* | 2B11+PD7/263 | Novus Biologicals | NBP2-34528 | 56 | 1:200 | 167 | 46 |
| *CD38* | EPR4106 | Abcam | ab176886 | 3 | 1:200 | 333 | 47 |
| *CD44* | IM7 | Biolegend | 103002 | 45 | 1:200 | 250 | 48 |
| *CD15* | MMA | BD Biosciences | 559045 | 14 | 1:200 | 25 | 49 |
| *Lambda light chain* | Lamb14 | Cell Marque | custom | 26 | 1:200 | 118 | 50 |
| *Mast cell tryptase* | AA1 | Abcam | ab2378 | 71 | 1:200 | 250 | 51 |
| *DRAQ5* | n/a | Cell Signaling Technology | 4084L | n/a | 1:100 | 167 | 52 |
| *Hoechst 33342* | n/a | Thermo Fisher Scientific | 62249 | n/a | 1:600 | 7 | all cycles |

**Supplemental Table 5: B-cell maturation state markers**

| Marker* | Maturation State(s) | | Physiological Function(s)** | CITE-Seq Panel | Reference*** |
|---|---|---|---|---|---|
| *IGKC (Kappa)* | All | Clonality indicator | Antigen and Ig receptor binding, immune activation, phagocytosis, Ig immune response to other organisms | ✅ | *Seifert M et al, Methods Mol Biol, 2019* |
| *IGLCx (Lambda)* | All | Clonality indicator | Antigen and Ig receptor binding, immune activation, phagocytosis, Ig immune response to other organisms | ✅ | *Seifert M et al, Methods Mol Biol, 2019* |
| *CXCR3* | Plasma<br>Memory | | GPCR for CXCL9, CXCL10 and CXCL11 for leukocyte traffic and chemotactic migration. | ❌ | *Morgan D and Tergaonkar V, Trends in Immunology, 2022.* |
| *IGHE* | Centrocyte (LZ)<br>Plasma<br>Memory | Late (class-switched) | Antigen and Ig receptor binding, immune activation, phagocytosis, Ig immune response to other organisms, circulating Ig complex | ❌ | *Talay O et al, Nature immunology, 2012* |
| *IGHAx* | Centrocyte (LZ)<br>Plasma<br>Memory | Late (class-switched) | Ig receptor binding, antibacterial humoral response, glomerular filtration, respiratory burst. | ❌ | *Morgan D and Tergaonkar V, Trends in Immunology, 2022.* |
| *IGHGx* | Centrocyte (LZ)<br>Plasma<br>Memory | Late (class-switched) | Antigen and Ig receptor binding, immune activation, phagocytosis, Ig immune response. | ❌ | *Morgan D and Tergaonkar V, Trends in Immunology, 2022.* |
| *FAS (CD95)* | (Pre)Memory | | TNF-receptor, pro-apoptotic regulator, activation of NF-kappaB, MAPK3/ERK1 and MAPK8/JNK. | ✅ | *Laidlaw B et al, Nat. Immunol., 2020* |
| *CD38* | Centrocyte<br>Plasma<br>Memory | Late (activated) | Synthesis and hydrolysis of cADP for intracellular calcium flux. | ✅ | *Morgan D and Tergaonkar V, Trends in Immunology, 2022.* |
| *PRDM1* | Plasma(blasts) | | Repressor of beta-interferon gene expression | ❌ | *Holmes A et al, J. Exp. Med., 2020* |
| *IRF4* | Plasma(blasts) | | Regulation of interferons and interferon-inducible genes, negatively regulates Toll-like-receptor (TLR) signaling. | ❌ | *Holmes A et al, J. Exp. Med., 2020* |

| | | | | | |
|---|---|---|---|---|---|
| *TNFRSF17* | Plasmablasts | | B-cell development, autoimmune response, NF-kappaB and MAPK8/JNK activation (via TNFSF13B/TALL-1/BAFF) . cell survival and proliferation (via TRAF) | ❌ | *Holmes A et al, J. Exp. Med., 2020* |
| *CCR6* | (Pre)Memory | | Beta chemokine receptor family, B-lineage maturation and antigen-driven B-cell differentiation, migration and recruitment of dendritic and T cells during inflammatory and immunological responses. | ❌ | *Morgan D and Tergaonkar V, Trends in Immunology, 2022.* |
| *BACH2* | Centrocytes (LZ) Memory | | Enables DNA binding, primary adaptive immune response | ❌ | *Sidwell et al, Nature, 2016* |
| *GPR183 (EBI2)* | (Pre)Memory | Early differentiation | Homing to outer B-cell follicle | ❌ | *Holmes A et al, J. Exp. Med., 2020* |
| *RELB* | Centrocytes (LZ) | Late | RNA polymerase and protein kinase binding. Lymphocyte differentiation and suppression of IF-beta production | ❌ | *Morgan D and Tergaonkar V, Trends in Immunology, 2022.* |
| *REL* | Centrocytes (LZ) | Late | Regulate genes involved in apoptosis, inflammation, the immune response, and oncogenic processes. B-cell survival and proliferation | ❌ | *Morgan D and Tergaonkar V, Trends in Immunology, 2022.* |
| *ICAM1* | Centrocytes (LZ) | Late | Binds CD11a and CD11b integrins, migration to endothelial tissue | ❌ | *Morgan D and Tergaonkar V, Trends in Immunology, 2022.* |
| *TRAF1* | Centrocytes (LZ) | Late | TNF receptor, NFKB activation, anti-apoptotic signalling | ❌ | *Morgan D and Tergaonkar V, Trends in Immunology, 2022.* |
| *NFKB2* | Centrocytes (LZ) | Late | Transcriptional regulation, inflammation, alteration of cell growth | ❌ | *Morgan D and Tergaonkar V, Trends in Immunology, 2022.* |

| | | | | | |
|---|---|---|---|---|---|
| *NFKB1* | Centrocytes (LZ) | Late | Transcriptional regulation, inflammation, alteration of cell growth | ❌ | *Morgan D and Tergaonkar V, Trends in Immunology, 2022.* |
| *CD40* | Centrocytes (LZ) | Late | TNF receptor, class-switching, memory B-cell development, germinal center formation | ❌ | *Morgan D and Tergaonkar V, Trends in Immunology, 2022.* |
| *BLNK* | Centrocytes (LZ) | | B-cell development | ❌ | *Morgan D and Tergaonkar V, Trends in Immunology, 2022.* |
| *CD74* | Centrocytes (LZ) | | Regulation of antigen presentation | ❌ | *Morgan D and Tergaonkar V, Trends in Immunology, 2022.* |
| *BLK* | Centrocytes (LZ) | | Cell proliferation, differentiation, B-cell receptor signalling, B-cell development | ❌ | *Morgan D and Tergaonkar V, Trends in Immunology, 2022.* |
| *BTK* | Centrocytes (LZ) | | B-cell development | ❌ | *Morgan D and Tergaonkar V, Trends in Immunology, 2022.* |
| *EBI3* | Centrocytes (LZ) | | Formation of IL-27 (regulation of T cells) | ❌ | *Morgan D and Tergaonkar V, Trends in Immunology, 2022.* |
| *CD83* | Centrocytes (LZ) | | Regulation of antigen presentation | ❌ | *Morgan D and Tergaonkar V, Trends in Immunology, 2022.* |
| *FOXP1* | LZ -> DZ | Transition | Transcription regulation (unspecified) | ❌ | *Morgan D and Tergaonkar V, Trends in Immunology, 2022.* |
| *CFLAR* | LZ -> DZ | Transition | Regulation of apoptosis | ❌ | *Morgan D and Tergaonkar V, Trends in Immunology, 2022.* |

| | | | | | |
|---|---|---|---|---|---|
| *FCRL2* | LZ -> DZ | Transition | Ig receptor | ❌ | *Morgan D and Tergaonkar V, Trends in Immunology, 2022.* |
| *SLA* | LZ -> DZ | Transition | Cell differentiation; innate immune response; and transmembrane receptor protein tyrosine kinase signaling pathway | ❌ | *Morgan D and Tergaonkar V, Trends in Immunology, 2022.* |
| *PTPN6* | DZ -> LZ | Transition | Hematopoietic cell growth, differentiation, mitotic cycle, and oncogenic transformation | ❌ | *Morgan D and Tergaonkar V, Trends in Immunology, 2022.* |
| *MS4A1 (CD20)* | DZ -> LZ | Transition | B-cell differentiation | ✅ | *Morgan D and Tergaonkar V, Trends in Immunology, 2022.* |
| *CD72* | DZ -> LZ | Transition | Enable signaling receptor binding activity, cell adhesion | ❌ | *Morgan D and Tergaonkar V, Trends in Immunology, 2022.* |
| *CAMK1* | DZ -> LZ | Transition | Calmodulin-dependent protein kinase cascade | ❌ | *Morgan D and Tergaonkar V, Trends in Immunology, 2022.* |
| *AICDA* | Centroblasts (DZ) | | Somatic hypermutation | ❌ | *Morgan D and Tergaonkar V, Trends in Immunology, 2022.* |
| *CCNB1* | Centroblasts (DZ) | Proliferating | Mitosis | ❌ | *Cattoretti G et al, Blood, 2006* |
| *BCL6* | Centroblasts (DZ) | | Transcription repression, blocking IL-4 response | ❌ | *Morgan D and Tergaonkar V, Trends in Immunology, 2022.* |
| *MME (CD10)* | Germinal centre (DZ.LZ) | Mainly centroblasts | Endopeptidase, hormone inactivation | ✅ | *Goteri G et al, Diagn. Pathol., 2017.* |
| *SELL (CD62L)* | Naïve | | Lymph node homing | ✅ | *Morgan D and Tergaonkar V, Trends in* |

| | | | | |
|---|---|---|---|---|
| | | | | *Immunology, 2022.* |
| *TCL1A* | Naïve | Cell survival (activation of AKT) | ❌ | *Morgan D and Tergaonkar V, Trends in Immunology, 2022.* |
| *IGHM* | Naïve<br>Memory | Antigen and Ig receptor binding, Ig secretion | ❌ | *Morgan D and Tergaonkar V, Trends in Immunology, 2022.* |
| *IGHD* | Naïve | Antigen and Ig receptor binding, IL-1 stimulation | ❌ | *Morgan D and Tergaonkar V, Trends in Immunology, 2022.* |

* Marker names are obtained from the NCBI Gene Database.  Aliases are listed in brackets if used elsewhere in this study.

** Gene functions from the NCBI Gene database are presented in summary form.

*** Publication linking the marker with its respective matur

**Supplemental Table 6: Full sample and library overview**

**Supplemental Table 7: List of target genes and chromosomal positions profiled with DNA sequencing**

**Supplemental Table 8: QC metrics for targeted DNA sequencing**

### Figures

#### Supplemental Fig. 1: FACS and classification of B-cell maturation states in reactive lymph nodes

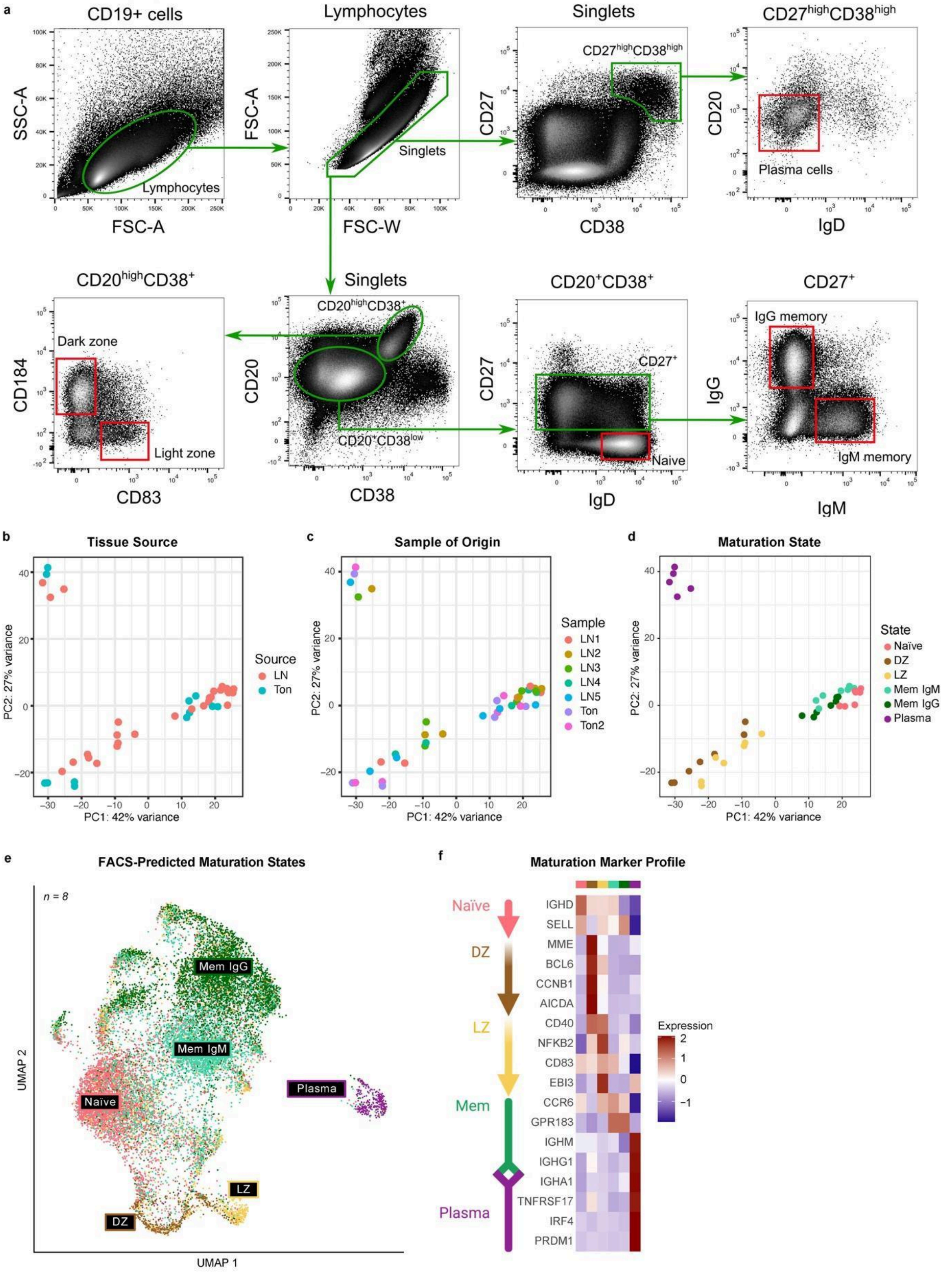

**a**, B-cell maturation state gating strategy for flow-activated cell sorting (FACS) employed on B cells isolated from human rLN (n=5) and tonsils (n=2) using the following marker panels: Naïve B cells (CD19+, CD20+, CD38low, CD27-, IgD high), germinal center dark zone B cells (CD19+, CD20+, CD38+, CD184+, CD83-), germinal center light zone B cells (CD19+, CD20+, CD38+, CD184-, CD83+), IgM memory B cells (CD19+, CD20+, CD38 low, CD27+, IgM+), IgG memory B cells (CD19+, CD20+, CD38 low, CD27+, IgG+), and plasmablasts/plasma cells (CD19+, CD20 low, CD38 high, CD27 high, IgD low). **b-d**, The first two principal components of RNA-seq data from FACS-sorted B-cell maturation states from rLN (n = 5) and tonsils (n = 2) colored by (b) tissue source, (c) the sample of origin, and (d) maturation state. **e**, Transcriptomic UMAP of the integrated CITE-Seq B cells data from 8 rLN labeled by maturation state predicted with logistic regression from the sorted states' RNA-seq data. **f**, Z-scaled gene expression of a subset of B-cell maturation markers (rows) across predicted maturation states (columns) in the rLN reference.

#### Supplemental Fig. 2: Isolation of malignant B-cells based on light chain restriction

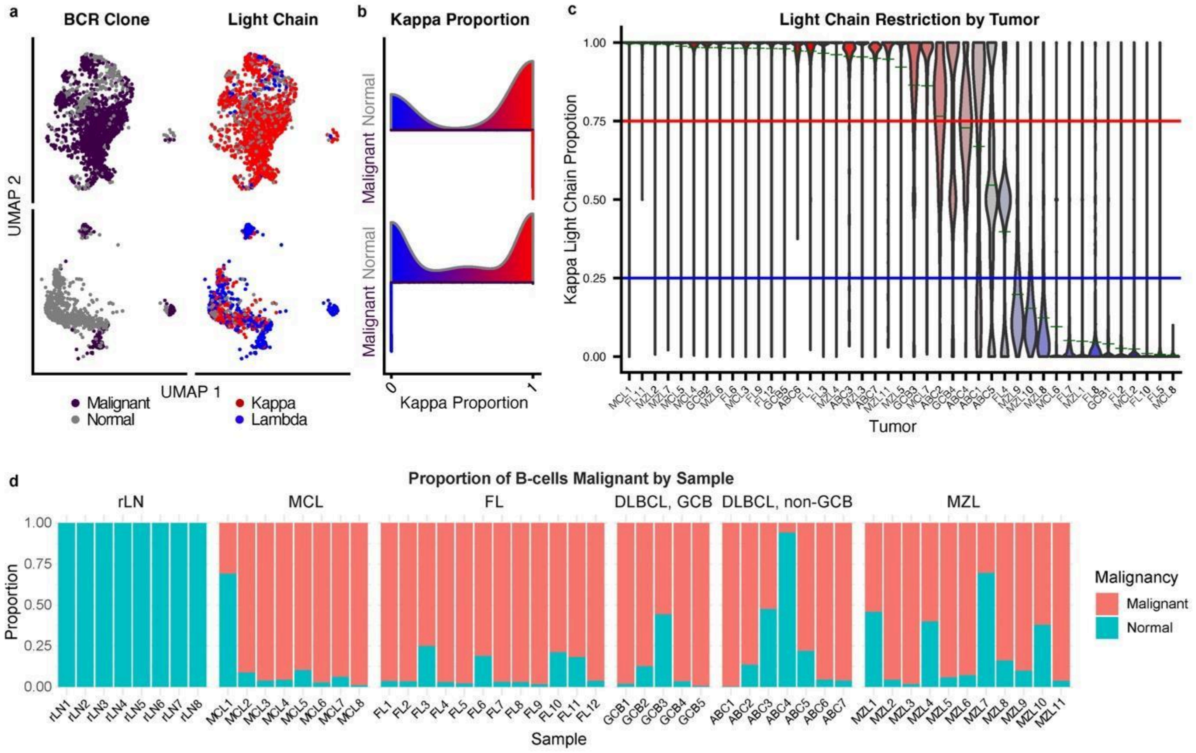

**a**, Reference-based UMAP labeled by malignant clone as determined by BCR profiling (left) and immunoglobulin light chain (right) for a MCL (top) and MZL (bottom) sample. **b**, Horizontal violin plot depicting the proportion of kappa light chain gene expression (x-axis) in malignant and normal B cells in the samples shown in (a). Red = kappa-positive, blue = lambda-positive. **c**, Vertical violin plot showing the proportion of kappa light chain surface epitope detected in malignant cells isolated from all tumor samples in the CITE-Seq cohort (n = 43). Red = kappa-restricted, blue = lambda-restricted. Malignant B cells were identified as light chain-restricted transcriptional clusters (mean kappa proportion >0.75 or <0.25). A light-chain-restricted tumor population was identified in all samples except ABC5 and FL4, which showed light-chain depletion instead. **d**, The proportion of B cells that are malignant or non-malignant, based on light chain restriction, in each sample, faceted by entity: reactive lymph nodes (rLN), mantle cell lymphoma (MCL), follicular lymphoma (FL), germinal center and non-germinal center diffuse large B-cell lymphoma (DLBCL, GCB/non-GCB), and marginal zone lymphoma (MZL).

#### Supplemental Fig. 3: B-cell maturation marker expression and maturation gene signature scores by entity

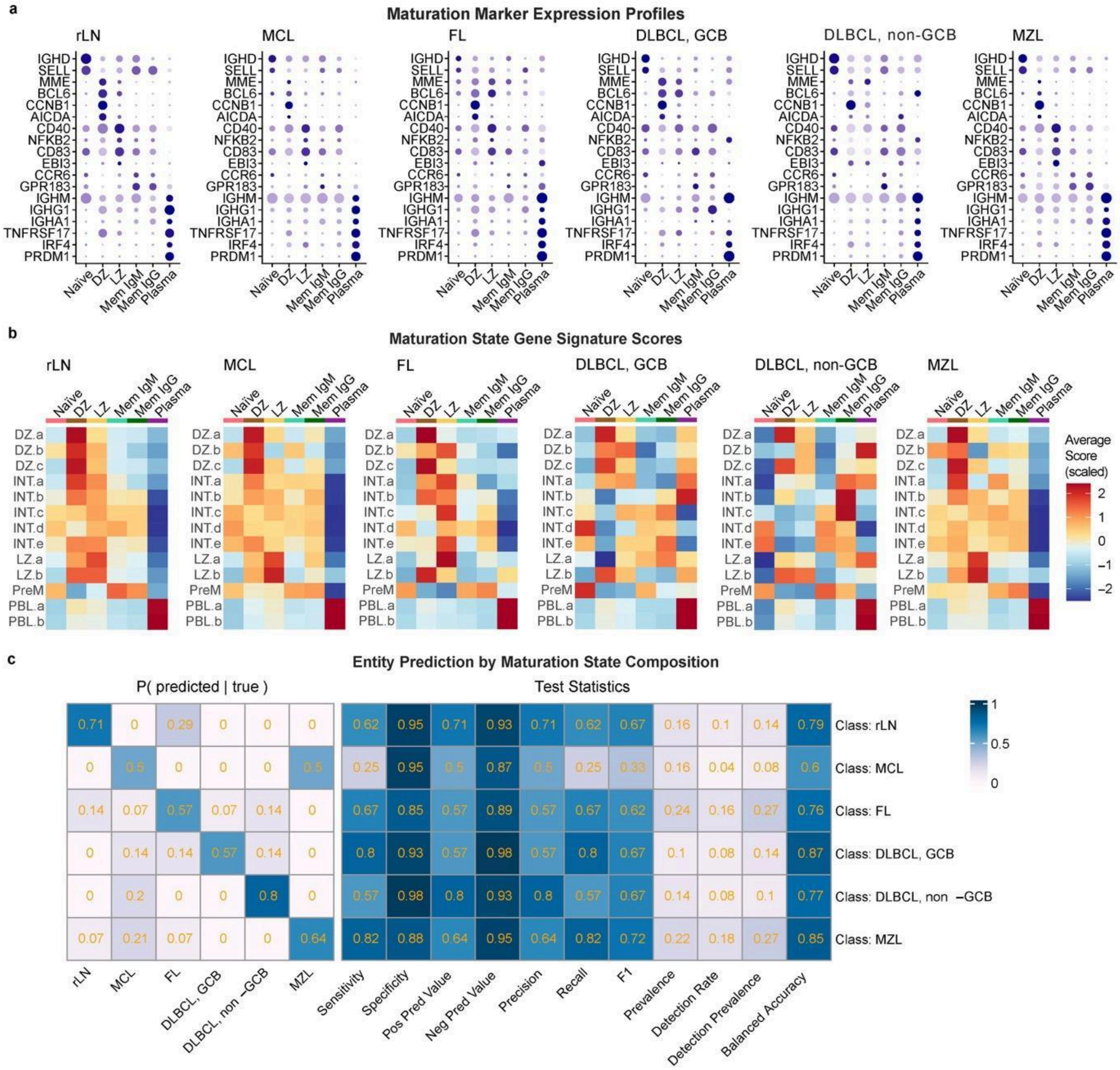

**a**, Dot plots showing the relative gene expression and abundance of maturation markers for each predicted maturation state in each entity. **b**, Heatmap of maturation state scores calculated from annotated maturation states in a published tonsil germinal center scRNA-seq dataset[2]. Each score shown is scaled across all scores in the dataset (mean = 0, sd = 1). **c**, Confusion matrix (left) showing the predicted (x) vs true (y) classes when predicting entity by maturation state proportions with random forest (10✕10-fold nested cross-validation), with test statistics (right) for classification of each entity (overall accuracy 63%). Entities: reactive lymph nodes (rLN), mantle cell lymphoma (MCL), follicular lymphoma (FL), germinal center and non-germinal center diffuse large B-cell lymphoma (DLBCL, GCB/non-GCB), and marginal zone lymphoma. See Fig. 1 for maturation state annotations.

#### Supplemental Fig. 4: Maturation state composition of samples

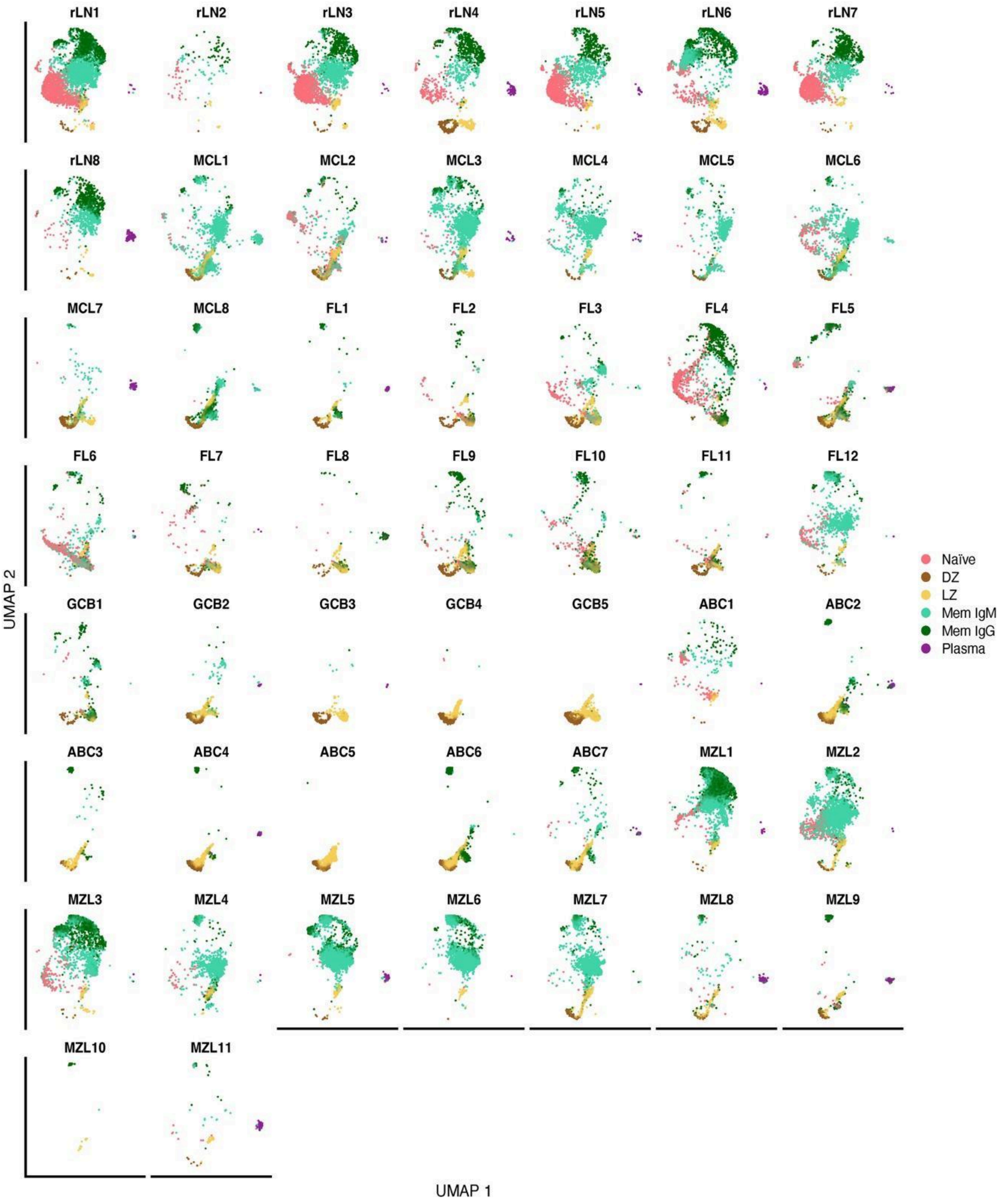

Reference-based UMAP labeled by B-cell maturation states for the CITE-Seq data from each sample (n=51). Maturation states are assigned by label transfer from the reactive lymph node reference in Fig. 1 as outlined in the Methods. In tumor samples, malignant cells were isolated from non-malignant B cells based on light chain restriction of transcriptional clusters (Supplemental Fig. 2). Maturation state annotations: Naïve = Naïve B cells, DZ = Centroblasts from the dark zone of the germinal center, LZ = Centrocytes from the light zone of the germinal center, Mem IgM = IgD+ and IgM+ memory B cells, Mem IgG = class-switched (IgG+ or IgA+) memory B cells, Plasma = plasma cells.

#### Supplemental Fig. 5: Multimodal subpopulation mapping of nodal B-cell non-Hodgkin lymphomas

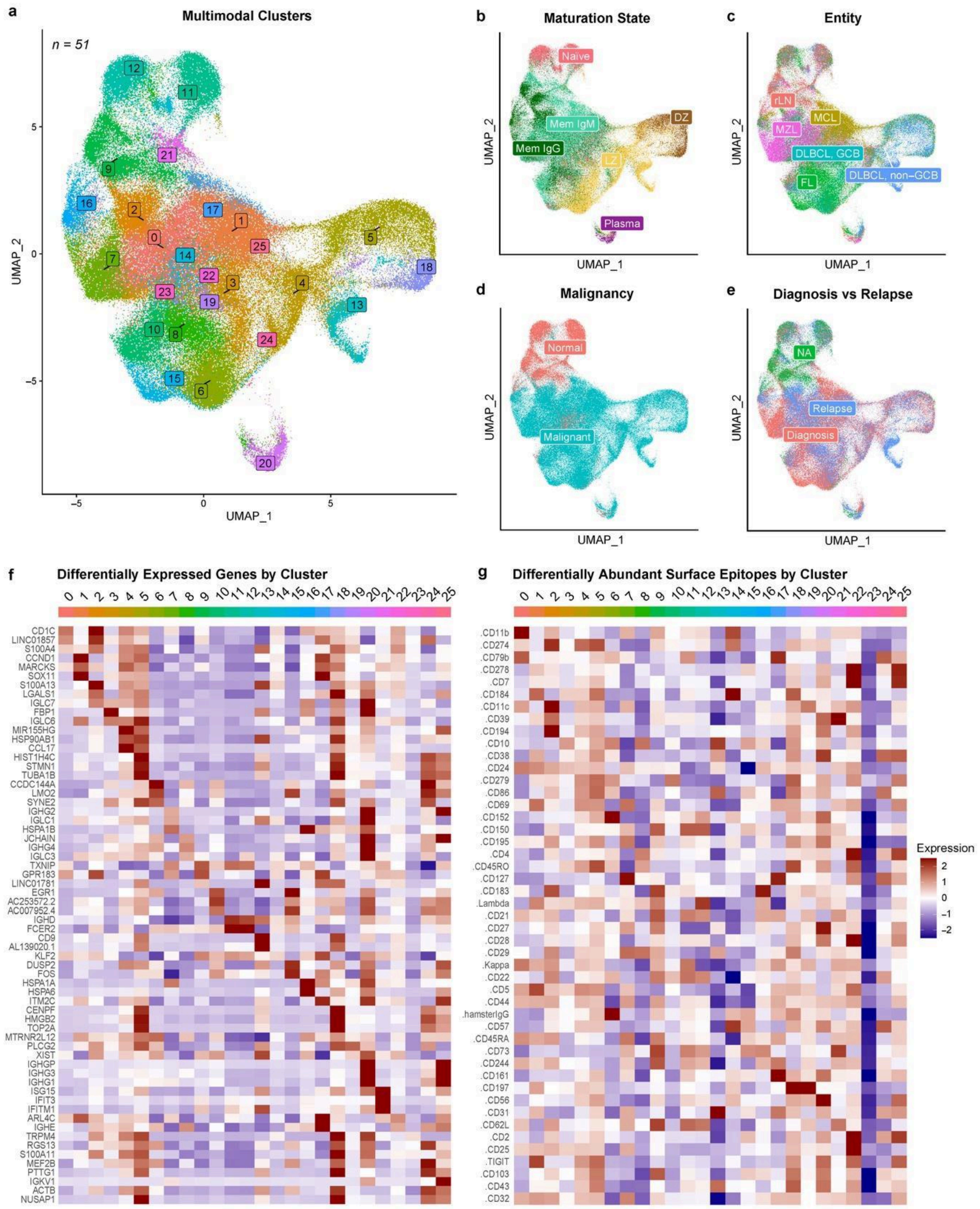

UMAP visualization of the full CITE-Seq B cells dataset (n=51) constructed with the latent factors (n = 50) from multi-omic factor analysis (MOFA)[3] based integration of the single-cell RNA and ADT (surface markers) data layers as principle components, labeled by **a**, clustering on the multimodal latent factor space, **b**, maturation states mapped from the reactive lymph node reference (Fig. 1b), **c**, entity, **d**, malignancy as determined by light chain restriction (Supplemental Fig. 2c), and **e**, samples taken at diagnosis or relapse. Z-scaled expression across multi-modal clusters of the 3 most differentially expressed genes (**f**) and proteins (**g**) by fold-change per cluster.

#### Supplemental Fig. 6: Longitudinal patterns of tumor maturation state composition

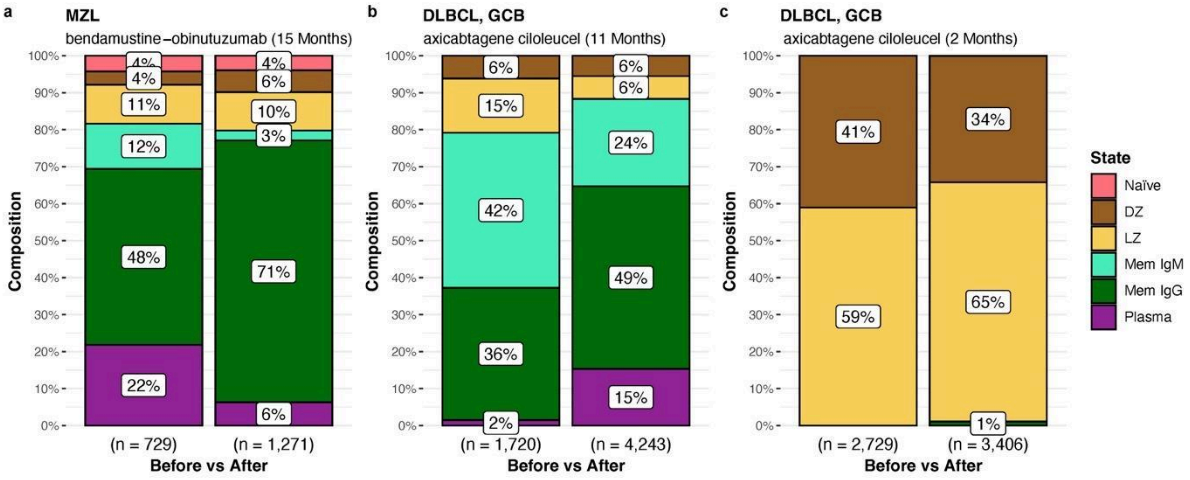

Maturation state composition of tumor cells in longitudinal samples from 3 patients: **a**, An MZL patient who relapsed 15 months following complete response to 6 cycles of obinutuzumab-bendamustine chemo-immunotherapy; **b-c**, two GCB DLBCL patients who relapsed after 11months (b) and 2 months (c) following axicabtagene ciloleucel (CAR-T cell) therapy. See Fig. 1 for maturation state annotations.

#### Supplemental Fig. 7: Differential transcription factor activity between tumor maturation states

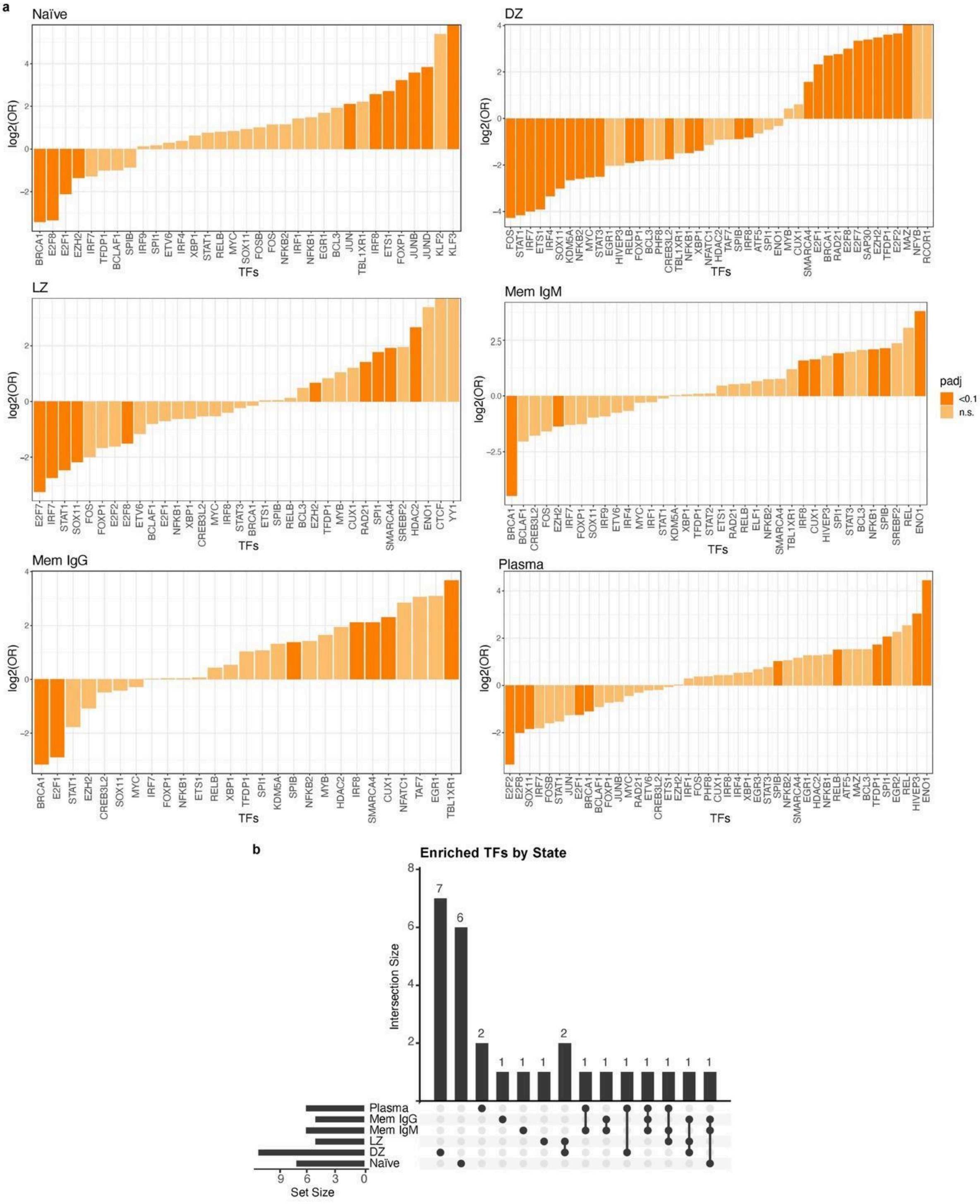

**a**, Bar charts showing the log2 odds ratio for differentially active transcription factors (TFs) in the tumor cells of each B-cell maturation state inferred with the *SCENIC* python package[4] from single-cell RNA-sequencing data from the malignant cells of all tumor samples combined (n=43). Only TFs with differentially expressed target genes (log2 fold-change >0.4, p < 10e-16 as determined with the *MAST* R package[5]) are shown. Bars highlighted in darker orange represent transcription factors with significant differential activity (FDR < 0.1

threshold). **b**, UpSet plot showing the intersections between the differentially active transcription factors (FDR <0.1) for each maturation state. See Fig. 1 for maturation state annotations.

#### Supplemental Fig. 8: Maturation state correlations between CITE-Seq and CODEX by sample

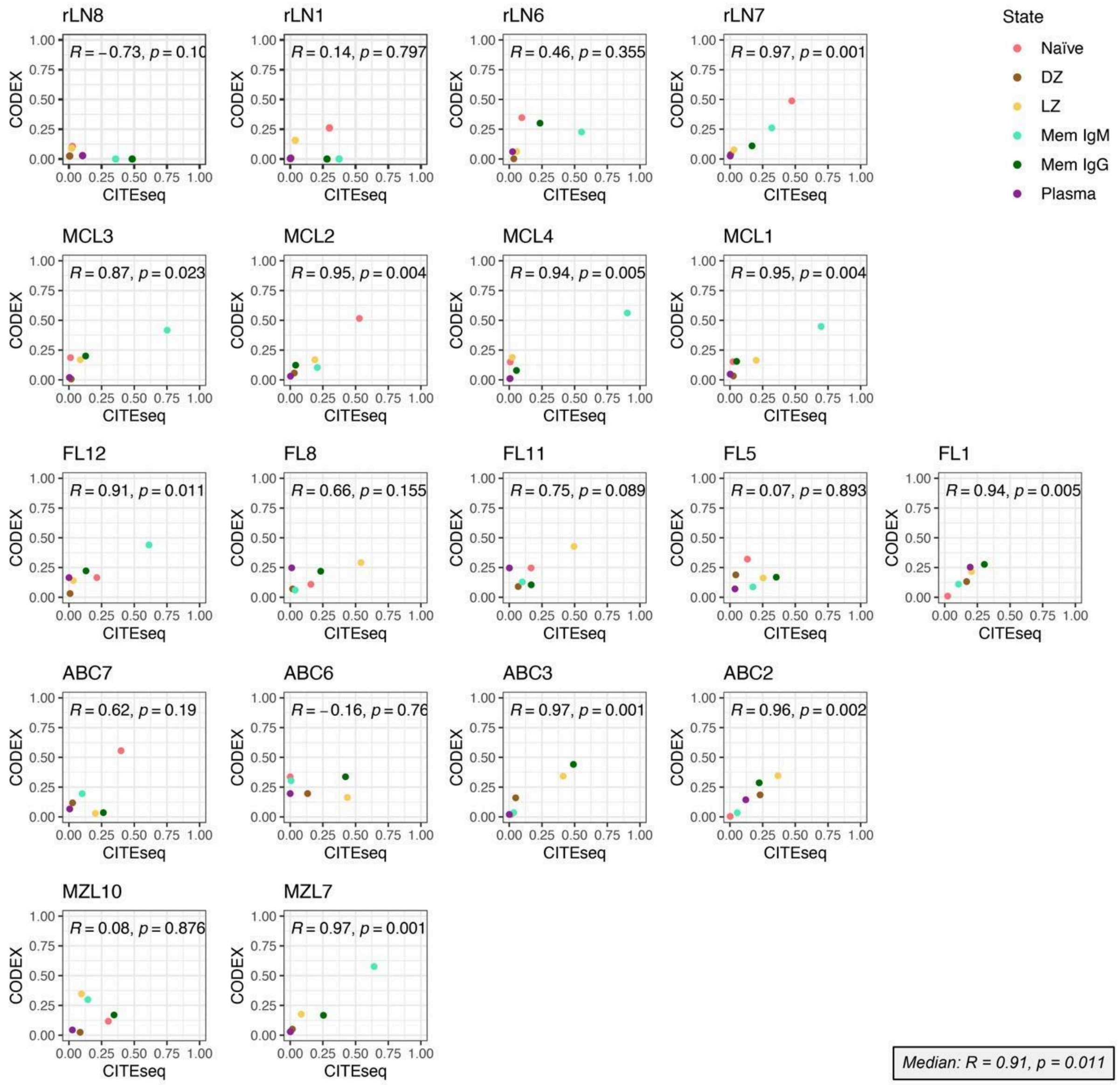

Scatter plots showing the Pearson correlation between B-cell maturation state proportions in the CITE-Seq and CODEX data for each **a**, reactive lymph node (rLN), **b**, mantle cell lymphoma (MCL), **c**, follicular lymphoma (FL), **d**, diffuse large B-cell lymphoma (DLBCL) and **e**, marginal zone lymphoma (MZL) sample. The median Pearson correlation coefficient across samples is shown (bottom-right). Maturation state labels were transferred from the CITE-Seq to CODEX data for each sample using logistic regression on shared protein channel/ADT features (n = 28). See Fig. 1 for maturation state annotations.

#### Supplemental Fig. 9: Spatial distribution of tumor microenvironments and maturation states

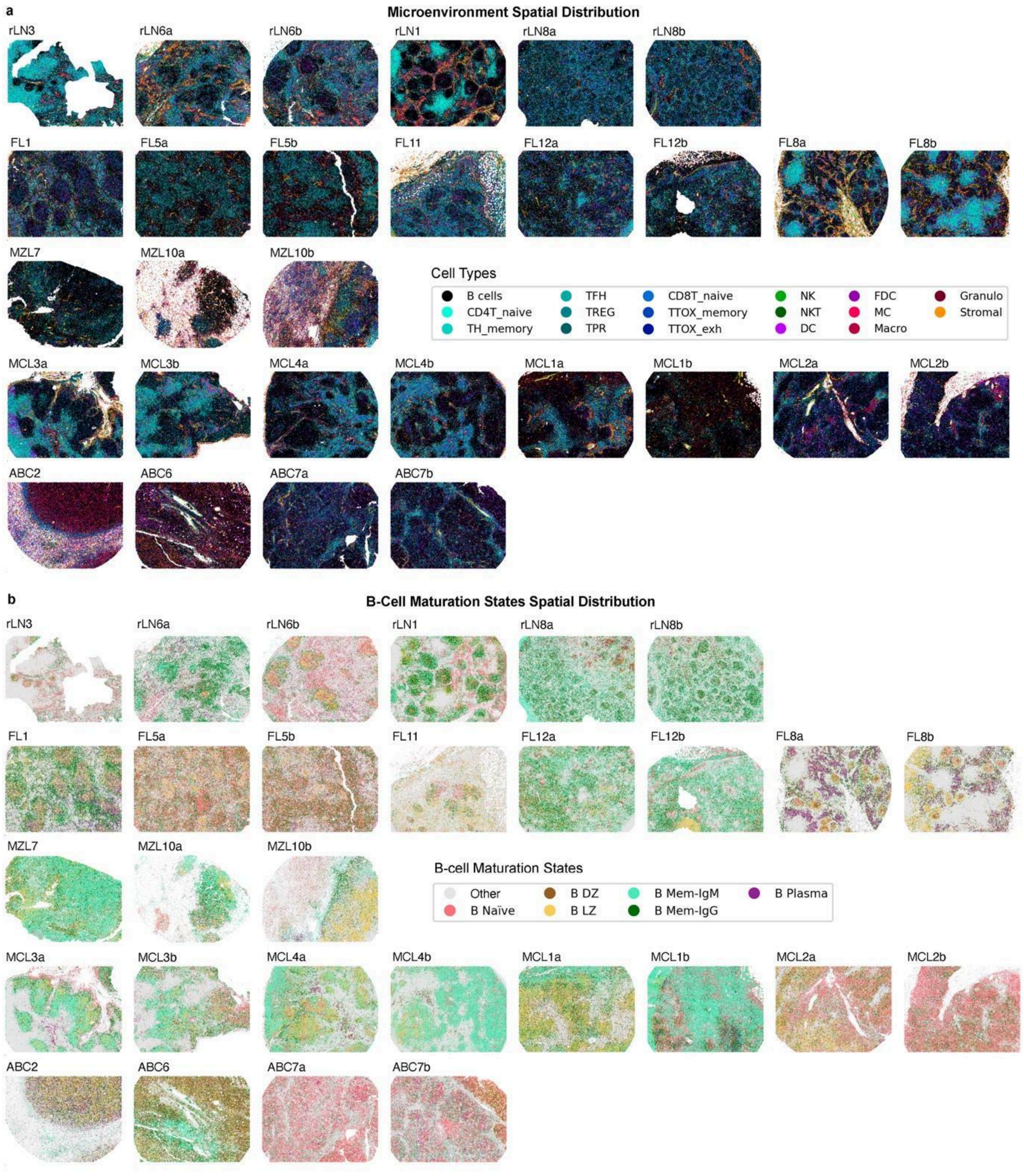

Spatial distribution of **a,** all cell types (with B cells in black) and **b,** B-cell subsets in reactive lymph nodes (rLN), mantle cell lymphoma (MCL), follicular lymphoma (FL), diffuse large B-cell lymphoma (DLBCL) and marginal zone lymphoma (MZL) from CODEX images on FFPE sections. B-cell states were classified samplewise using logistic regression from the CITE-Seq data using the shared features (n = 28). Marker-based annotation of all other cell types was performed on clustering of the CODEX features (n = 52, see Supplemental Table 5). B Naïve = naïve B cells, B DZ = centroblasts from the dark zone of the germinal center, B LZ = centrocytes from the light zone of the germinal center, B Mem IgM = IgD+ and IgM+ memory B cells, B Mem

IgG = class-switched (IgG+ or IgA+) memory B cells, B Plasma = plasma cells, CD4T_naive = naive CD4+ T-cells, CD8T_naive = naive CD8+ T-cells, TH_memory = memory helper T-cells, TTOX_memory = memory cytotoxic T-cells, TTOX_exh = exhausted cytotoxic T-cells, NKT = natural killer T-cells, TFH = follicular helper T-cells, TPR = proliferating T-cells, TREG = regulatory T-cells, FDC = follicular dendritic cells, DC = dendritic cells, Macro = macrophages, Stromal = stromal cells, NK = natural killer cells, MC = monocytes, Granulo = granulocytes.

#### Supplemental Fig. 10: Spatial distribution of cellular neighborhoods

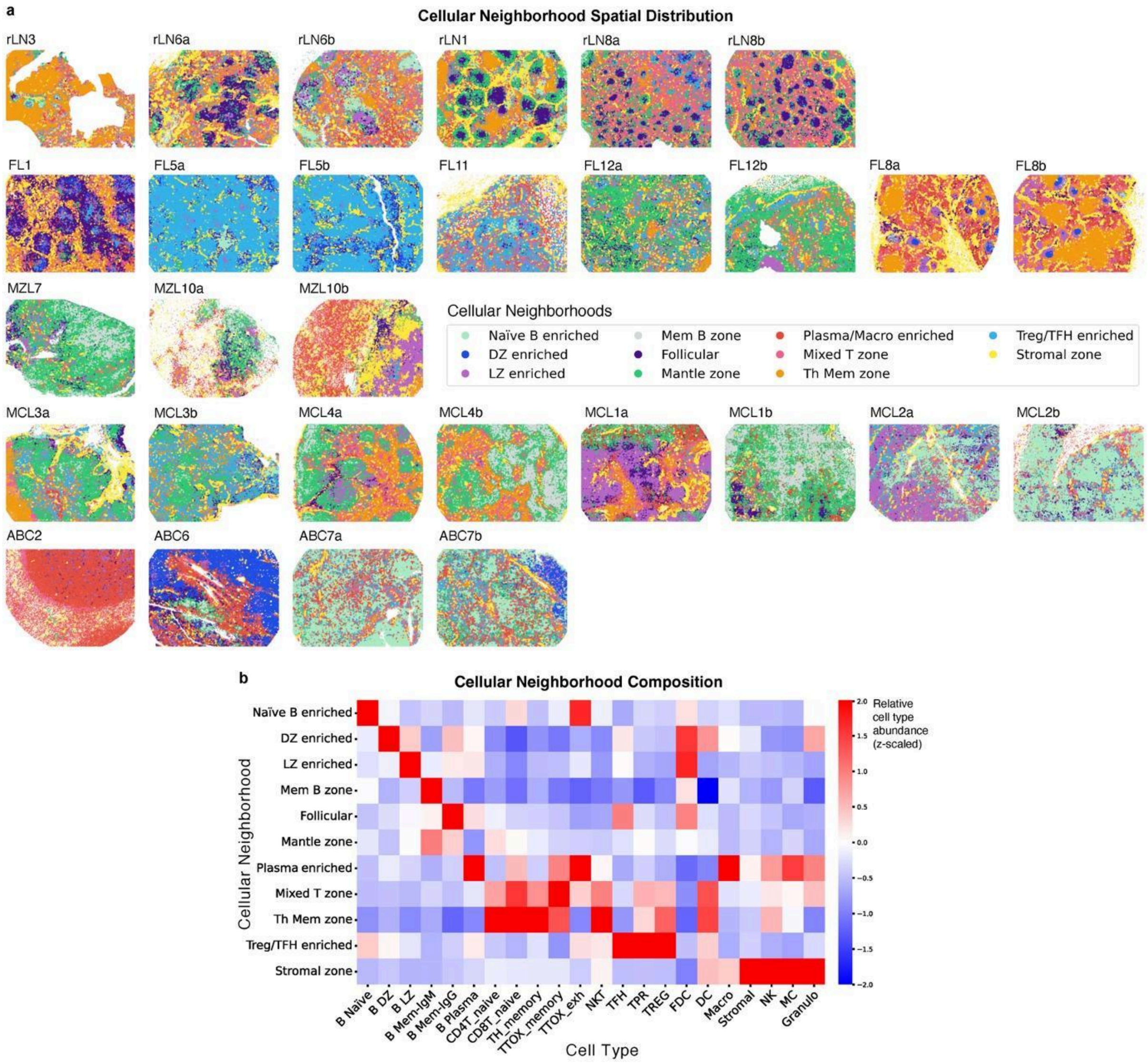

**a**, Spatial distribution of cellular neighborhoods (CNs) in reactive lymph nodes (rLN), mantle cell lymphoma (MCL), follicular lymphoma (FL), diffuse large B-cell lymphoma (DLBCL), and marginal zone lymphoma (MZL) from CODEX images on FFPE sections. CNs (n = 11, KNN = 20) were calculated using all CODEX slides (n=29) and labeled based on their distinguishing features, ie. tumor cells' predominant maturation state (eg. DZ, Mem), B cells' location or function in reactive lymph nodes (eg. Follicular, Mantle zone), or enriched cell type(s) (eg. Mixed T zone, T Mem zone). **b**, Relative abundance of each cell type across Cellular Neighborhoods (CNs), scaled by cell type frequency. B Naïve = naïve B cells, B DZ = centroblasts from the dark zone of the germinal center, B LZ = centrocytes from the light zone of the germinal center, B Mem IgM = IgD+ and IgM+ memory B cells, B Mem IgG = class-switched (IgG+ or IgA+) memory B cells, B Plasma = plasma cells, CD4T_naive = naive CD4+ T-cells, CD8T_naive = naive CD8+ T-cells, TH_memory = memory helper T-cells, TTOX_memory = memory cytotoxic T-cells, TTOX_exh = exhausted cytotoxic T-cells, NKT = natural killer T-cells, TFH = follicular helper T-cells, TPR = proliferating T-cells, TREG = regulatory T-cells, FDC = follicular dendritic cells, DC = dendritic cells, Macro = macrophages, Stromal = stromal cells, NK = natural killer cells, MC = monocytes, Granulo = granulocytes.

#### Supplemental Fig. 11: Mutation landscape of the overall cohort

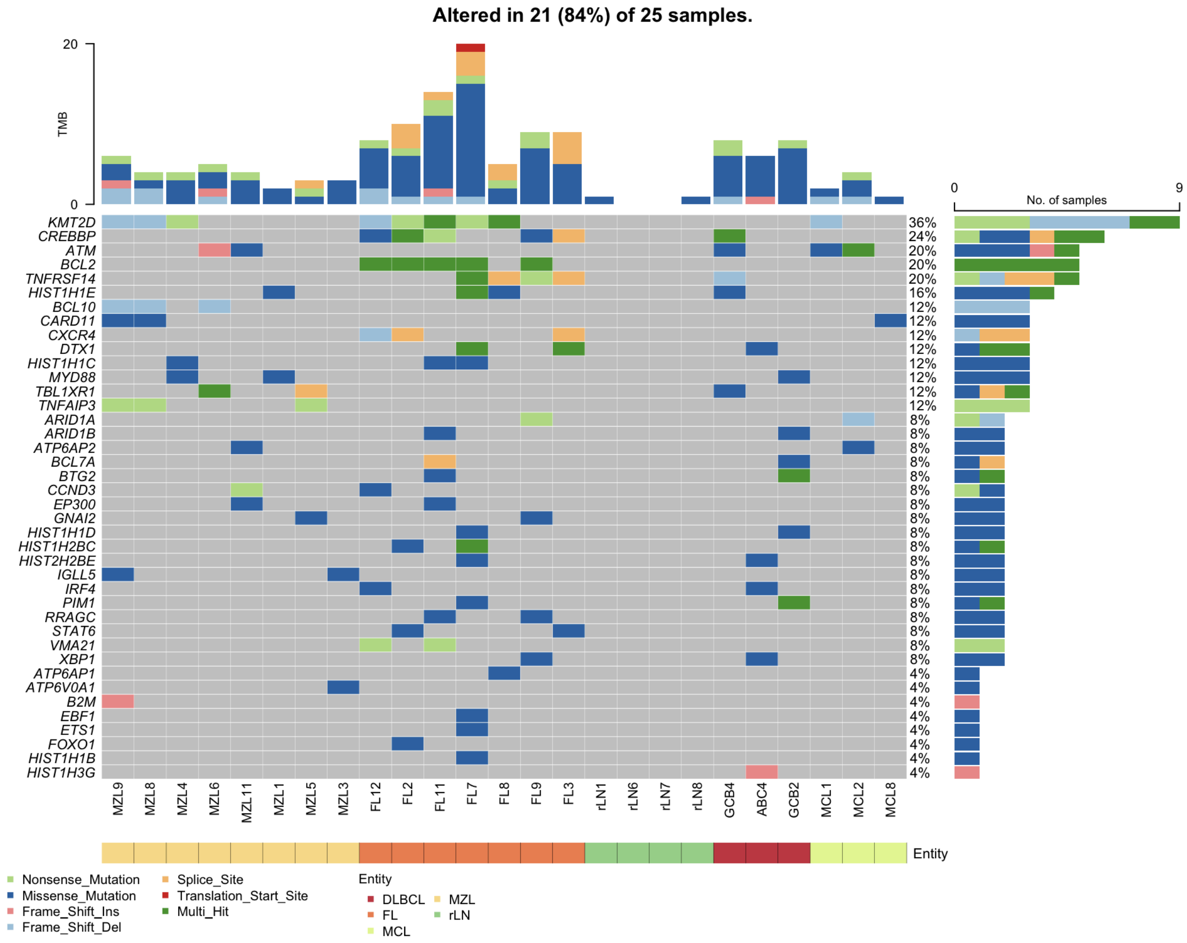

Mutation landscape of a subset of the CITE-Seq cohort determined with target DNA-sequencing (see targets in . All non-silent mutations with VAF >= 10% are depicted. Reactive lymph nodes (rLN), mantle cell lymphoma (MCL), follicular lymphoma (FL), diffuse large B-cell lymphoma (DLBCL), and marginal zone lymphoma (MZL).
